# Clair-Mosaic: A deep-learning method for long-read mosaic small variant calling

**DOI:** 10.1101/2025.10.31.685831

**Authors:** Lei Chen, Zhenxian Zheng, Minggao He, Angel On Ki Wong, Xian Yu, Junzhe Li, Jingcheng Zhang, Yekai Zhou, Ruibang Luo

## Abstract

Mosaic variants, defined as postzygotic mutations occurring during an organism’s development from zygote to adult, play critical roles in developmental biology, aging, and diseases such as cancer and neurological disorders. However, their accurate detection remains challenging due to low abundance in the genome and low variant allelic fractions (VAF). While current mosaic variant callers are primarily designed for short-read sequencing, no method is available for long-read sequencing, which can generate tens of kilobase-long reads to cover complex genomic regions inaccessible to short reads. To fill the gap, we present Clair-Mosaic, a deep-learning-based method for detecting mosaic small variants from long-read data. Clair-Mosaic was trained on hundreds of millions synthetic variants encompassing diverse read coverages and allelic fractions, enabling it to detect low-VAF mosaic variants with high sensitivity in paired-sample and single-sample modes. In addition to neural network prediction, Clair-Mosaic distinguishes genuine mosaic variants from sequencing artifacts by leveraging their inherent haplotype relationship in phased long reads. Furthermore, a Bayesian mosaic-germline discriminator is introduced to distinguish mosaic variants from germline variants. It also employs multiple post-calling filters, including a mosaic variant database and multiple germline population resources, to tag common germline and mosaic variants. Comprehensive benchmarking on synthetic datasets and real samples demonstrated Clair-Mosaic’s outstanding performance in ONT and PacBio. Clair-Mosaic is also applicable to short-read data and outperforms methods like MosaicHunter, MosaicForecast, DeepMosaic, and DeepSomatic. Clair-Mosaic is open-source and available at https://github.com/HKU-BAL/Clair-Mosaic.

## Introduction

Somatic mosaicism, arising from postzygotic mutations that manifest in subsets of an organism’s cells, is a key driver of genomic diversity and cellular evolution within an individual^1–3^. The analysis of mosaic mutations in non-disease samples is critical for understanding normal developmental processes, tissue aging, and the pathogenesis of various diseases, including cancer and neurological disorders^4–11^. Consequently, accurately characterizing these variants is essential. However, their detection poses substantial technical challenges: a specific mosaic mutation may exist only in a small clone of cells, leading to low variant allelic fraction (VAF) in bulk sequencing data^6^. This inherent low VAF, low abundance, combined with the difficulty in distinguishing them from germline variants and sequencing artifacts in low-confidence genomic regions, makes mosaic variant detection particularly challenging, especially in noisy long-read sequencing data.

Short-read sequencing, the foundation of most current variant detection methods, is fundamentally limited by its read length when resolving repetitive or complex genomic regions^12, 13^. These limitations frequently result in ambiguous alignments and incomplete variant calls in such areas. Long-read sequencing technologies, such as those developed by Oxford Nanopore Technologies (ONT) and Pacific Biosciences (PacBio), overcome this barrier by producing reads tens of kilobases in length, which provide unique mappability and comprehensive haplotype resolution^13–15^. The demonstrated superiority of long-read sequencing in germline variant calling within complex regions highlights its potential for detecting mosaic variants^16–20^. Nevertheless, existing mosaic variant callers remain tailored to short-read data^21–24^, with none effectively optimized for long-read sequencing, leaving a notable methodological gap in the field.

A further challenge lies in the scarcity of well-characterized mosaic variants, which severely limits the method development and performance evaluation^24, 25^. For instance, while the seven Genome in a Bottle (GIAB) reference samples (HG001–HG007) provide ~32 million high-confidence germline variants^26^, the landscape of validated mosaic variants remains markedly underdeveloped. Currently, the only available benchmark derived from the GIAB HG002 sample contains merely 85 high-confidence mosaic SNVs^25^. And even in a non-disease individual, the number of mosaic variants typically ranges from only a few hundred to several thousand, a number far from robust model training. This pronounced scarcity of mosaic variants underscores the critical need for alternative strategies, such as the use of synthetic mosaic datasets, to facilitate reliable model development and performance assessment.

To address these limitations, we present Clair-Mosaic, the first publicly available deep-learning-based method specifically designed for long-read mosaic small variant calling. Clair-Mosaic offers dual modes, a paired-sample mode and single-sample mode, to provide flexibility for diverse research and clinical scenarios where matched samples may be unavailable. To overcome the scarcity of real mosaic variants for model training, Clair-Mosaic employs a synthetic data generation strategy. By combining sequencing reads from two distinct samples *in silico*, germline variants unique to one sample are considered as mosaic variants in a synthetic mixture. This approach enables the generation of training variants at a hundreds of millions scale, thereby providing the volume and diversity of training samples with diverse read coverages, allelic fractions, and genomic contexts. Clair-Mosaic also leverages phasing information for haplotype consistency checking, to identify mosaic variants with low VAF and incorporates multiple post filters to minimize false positives. Moreover, we developed a Bayesian Mosaic-Germline Discriminator (BayMGD) that integrates observed individual allelic fraction, population allele frequency and various quality metrics into a unified probabilistic framework to compute likelihood scores for competing mosaic versus germline hypotheses, effectively distinguishing true mosaic variants from germline polymorphisms.

We established a synthetic mosaic variant generation and benchmarking workflow using the GIAB Ashkenazi (HG002-HG004) and Chinese (HG005-HG007) trios^13, 26^ for comprehensive performance evaluation. When evaluated on these synthetic samples, Clair-Mosaic achieved F1-scores of 93.86% and 98.80% on ONT and PacBio HG002/HG003 synthetic datasets, respectively, consistently outperforming DeepSomatic^27^. On the set of 85 real mosaic variants in the GIAB HG002 sample, Clair-Mosaic detected 77 and 82 variants in the ONT and PacBio datasets, respectively. Moreover, Clair-Mosaic detected thousands of potential mosaic variants in complex genomic regions from both ONT and PacBio data that were absent in Illumina. Although designed for long-read data, the method is also applicable to Illumina short reads. This versatility enabled a benchmark on short-read data, where Clair-Mosaic surpassed leading short-read callers, including MosaicHunter^21^, MosaicForecast^22^, DeepMosaic^23^, and DeepSomatic^27^, on the HG002 sample. Across experiments with various sequencing platforms, read coverages, allelic fractions, and genomic contexts, we demonstrated that Clair-Mosaic is a reliable mosaic variant caller.

## Results

### Overview of Clair-Mosaic

An overview of the Clair-Mosaic variant calling workflow is presented in **Figure 1b**. Clair-Mosaic is a deep-learning-based mosaic small variant caller primarily designed for long-read sequencing data. It supports both paired-sample and single-sample analyses, allowing for application in scenarios with or without a matched control. In paired-sample mode, a pileup model processes the input and control samples by vertically concatenating the features to estimate the probability that a candidate variant is a mosaic variant, a germline variant, or a sequencing artifact. When only the input sample is available (single-sample mode), Clair-Mosaic employs a pileup model to classify each candidate variant as a mosaic or an artifact. Furthermore, Clair-Mosaic leverages the advantages of long-read data to perform haplotype consistency checking for low-VAF mosaic variants. Specifically, a variant candidate identified in only one haplotype is considered reliable, while a candidate appearing in both haplotypes is flagged as a potential artifact. To enhance specificity, Clair-Mosaic applies multiple germline resources to tag non-mosaic variants. Complementing these filters, Clair-Mosaic designs a Bayesian Mosaic-Germline Discriminator (BayMGD), a probabilistic framework that computes continuous posterior probabilities for mosaic and germline hypotheses by integrating observed individual allelic fraction, population frequency, and quality score metrics, thereby quantifying biological plausibility beyond hard thresholds.

**Figure 1.**
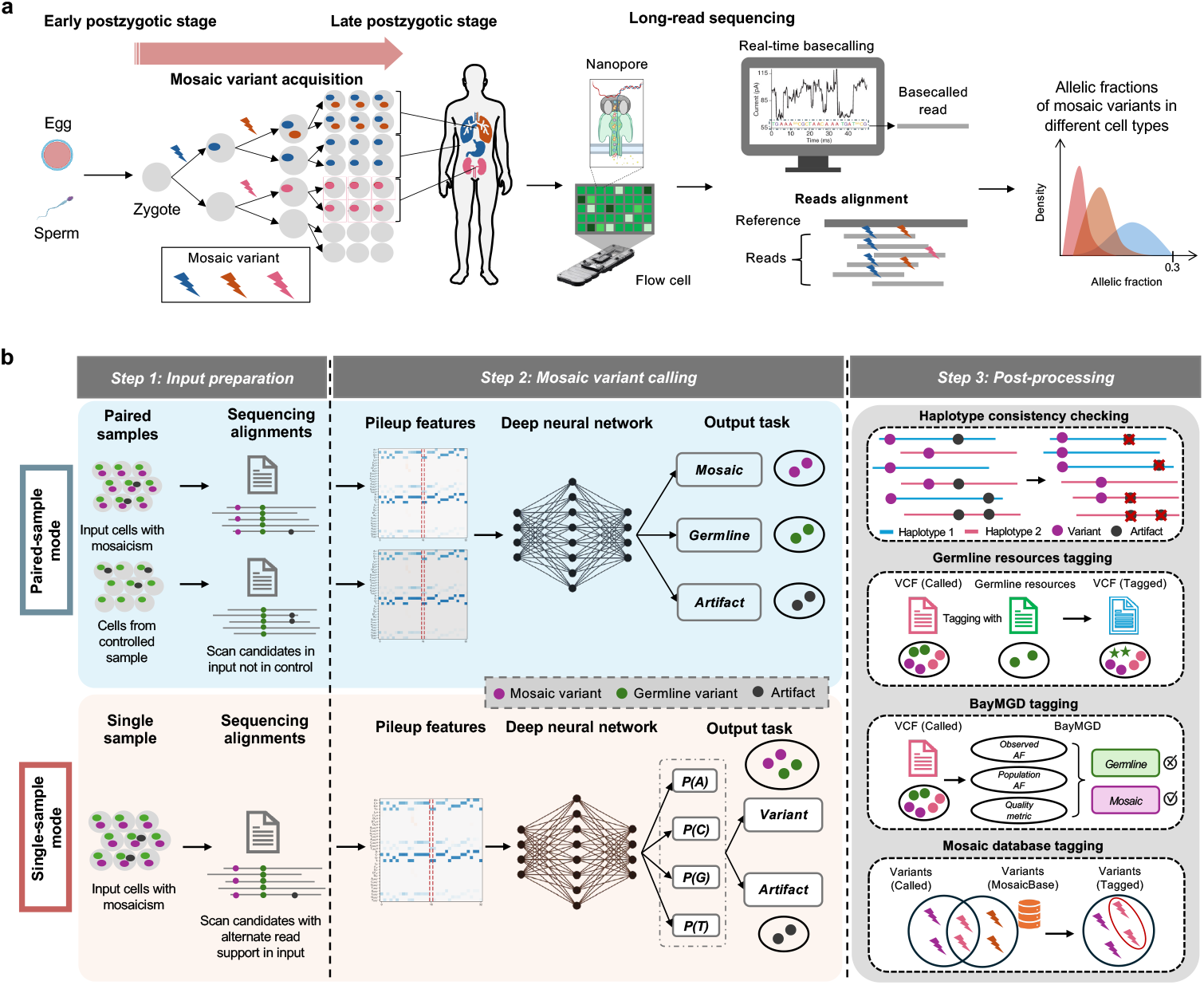
Overview of Clair-Mosaic variant calling workflow. **(a)** This figure illustrates the mechanism of postzygotic mosaicism. A variant originates in a single somatic cell and propagates through cell division, forming a clone of cells that all carry the mutation. Crucially, the variant allelic fraction observed in bulk sequencing data (e.g., from ONT) is determined by the developmental timing of the initial mutation. An early-arising variant undergoes more cell divisions, leading to a larger clone and a moderately high VAF. In contrast, a late-arising variant has limited clonal expansion, resulting in a smaller clone and a very low VAF in the sequenced tissue sample. **(b)** The figure illustrates the Clair-Mosaic variant calling workflow, which supports both paired-sample and single-sample analysis modes. In both configurations, sequencing data are first converted into pileup-level features for inference using a deep neural network. The model initially classifies candidate variants into mosaic, germline, or artifact categories. These preliminary calls subsequently undergo a series of post-filters—including haplotype consistency checking, germline resource (GR) tagging, and a Bayesian Mosaic-Germline Discriminator (BayMGD)—to accurately differentiate true mosaic variants from germline polymorphisms and technical artifacts.

### Benchmarking datasets

The extreme scarcity of validated benchmark sets for real mosaic variants severely restricts comprehensive method evaluation. To address this gap, we designed synthetic datasets to be used alongside real benchmark samples for evaluation. Synthetic datasets were generated by *in silico* mixing of reads from two biologically related samples, namely a child and one of their parents. This approach is based on the principle that postzygotic mosaic mutations are not inherited from parents. We constructed two independent sets of mixtures: one using the HG002/HG003/HG004 trio (comprising 724,280 SNVs and 151,226 Indels for the HG002/HG003 mixture, and 713,511 SNVs and 148,264 Indels for the HG002/HG004 mixture), and another using the HG005/HG006/HG007 trio (yielding 650,555 SNVs and 141,487 Indels for the HG005/HG006 mixture, and 639,590 SNVs and 141,557 Indels for the HG005/HG007 mixture), with the statistics shown in **Figure 2d**. These mixtures yield hundreds of thousands of synthetic mosaic variants, substantially exceeding the number of real mosaic variants for evaluation. As shown in **Figure 2a**, benchmarking regions were defined as the intersection of GIAB high-confidence regions between two samples, and truth variant sets were constructed by subtracting germline variants of the control sample from those of the input sample, thereby treating sample-specific germline variants as synthetic mosaics. In addition to the synthetic datasets, we also performed real-sample validation. Specifically, we utilized the GIAB HG002 mosaic variant benchmark v1.0 released by the GIAB Consortium^25^, which provides 85 high-confidence mosaic SNVs identified from a large batch of a normal lymphoblastoid cell line using multiple sequencing technologies, with the VAF distribution shown in **Figure 2c**. All benchmarks adhered to the same truth inclusion criteria: coverage ≥4, alternative allele depth ≥3, and VAF ≥0.05, to exclude variants without enough read evidence. Finally, we applied Clair-Mosaic to all seven GIAB samples (HG001– HG007) to provide a comprehensive assessment of mosaic variant detection across different individuals.

**Figure 2.**
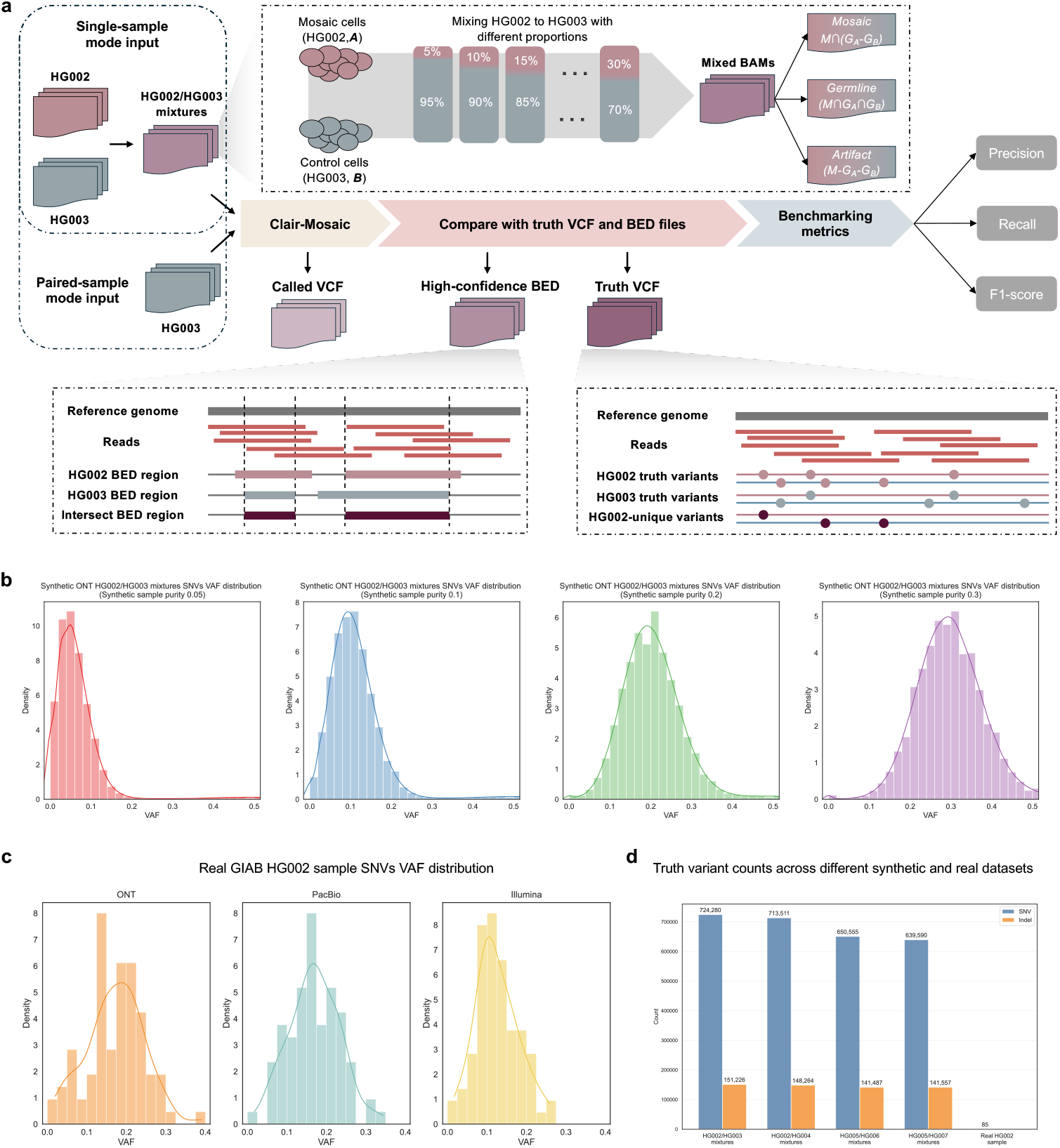
Synthetic mosaic variant benchmarking. **(a)** This figure illustrates the synthetic mosaic variant benchmarking workflow. The benchmark is constructed by mixing two GIAB samples (HG002 and HG003 as example) at varying purities to simulate mosaic events. Truth variants are defined as sample-specific germline variants (germline variants present in the input sample but absent in the control sample), which are treated as synthetic mosaic events. The high-confidence benchmarking region is defined as the intersection of GIAB high-confidence regions between the two samples. Performance metrics (precision, recall, F1-score) are obtained by comparing Clair-Mosaic calls against the constructed truth set within the defined high-confidence regions. **(b)** The figure illustrates the allelic fraction distribution in different sample purities on ONT HG002/HG003 synthetic dataset. **(c)** The figure illustrates the allelic fraction distribution in HG002 real mosaic variants in ONT, PacBio, and Illumina, respectively. **(d)** The synthetic SNV and Indel variants in different synthetic mixtures and real mosaic SNVs in the HG002 dataset.

### Performance on ONT data

A summary of the ONT datasets used for Clair-Mosaic model training and benchmarking is provided in **Supplementary Table 1**. For both paired-sample and single-sample modes, Clair-Mosaic adopted the training workflow of ClairS^28^ and ClairS-TO^26^, which are designed for somatic variant calling. The model was trained on synthetically mixed data generated from biologically independent HG002 and HG001 samples obtained from the ONT EPI2ME Labs^29^, to ensure generalizability by maximizing the training variants derived from two biologically unrelated samples. For benchmarking, we employed synthetic mixtures of biologically related samples from two independent family trios: HG002/HG003/HG004 and HG005/HG006/HG007. This approach aligns with the biological principle that postzygotic mosaic variants are not inherited from parents^1, 2^, thereby enabling the simulation of somatic mosaicism across diverse genomic contexts, VAF ranges, and read coverages. Due to the absence of dedicated long-read mosaic variant callers, we compared Clair-Mosaic against DeepSomatic, a deep learning-based somatic variant caller supporting both tumor–normal paired and tumor-only modes, which was trained on real cancer cell lines. For a fair comparison, we adhered to the authors’ recommended pipeline and parameter settings. Clair-Mosaic was not directly benchmarked against ClairS (optimized for tumor–normal somatic variant calling) and ClairS-TO (designed for tumor-only somatic variant calling), as the model training workflow and network architecture of Clair-Mosaic were adapted from the well-established designs of ClairS and ClairS-TO.

#### Performance at different sample purities

We evaluated Clair-Mosaic on synthetic datasets generated from two independent family trios: the Ashkenazi trio (HG002/HG003/HG004) and the Chinese trio (HG005/HG006/HG007), with results summarized in **Figure 3a, Supplementary Figure 1**, and **Supplementary Table 2**. To mimic the VAF distribution of real mosaic variants (typically 5–30%^1, 25^), we created mixtures at four sample purities (5%, 10%, 20%, and 30%) by *in-silico* mixing of sequencing reads from HG002 into HG003 (or HG004) and from HG005 into HG006 (or HG007), ensuring that the resulting VAF peaks aligned with the expected mosaic VAF range, with the VAF distribution shown in **Figure 2b**. In paired-sample mode, using 50-fold HG002 as input and 25-fold HG003 or HG004 as control for mixtures, Clair-Mosaic achieved F1-scores of 85.81%, 93.11%, 95.69%, and 96.38% at the four purity levels for the HG002/HG003 mixture, outperforming DeepSomatic by margins of 28.58%, 14.95%, 1.60%, and 0.60%, respectively. A similar performance advantage was observed for the HG002/HG004 mixture. In single-sample mode, Clair-Mosaic attained F1-scores of 82.68%, 91.70%, 93.32%, and 94.82%, while DeepSomatic scored only 23.31%, 26.68%, 29.34%, and 29.52%, resulting in performance gaps of 59.37%, 65.02%, 63.98%, and 65.30%, respectively. The performance gap is more pronounced in single-sample mode, primarily because DeepSomatic incorporates germline population frequency information directly into its neural network architecture. This design allows the model to more easily classify synthetic mosaic variants—which are, in reality, sample-specific germline polymorphisms—as non-mosaic events. Hence, the performance of DeepSomatic single-sample mode in our synthetic datasets is for reference only. Consistently, evaluation on the 50-fold HG005 as input and 25-fold HG006 or HG007 as control confirmed Clair-Mosaic’s robust performance across diverse genomic backgrounds.

**Figure 3.**
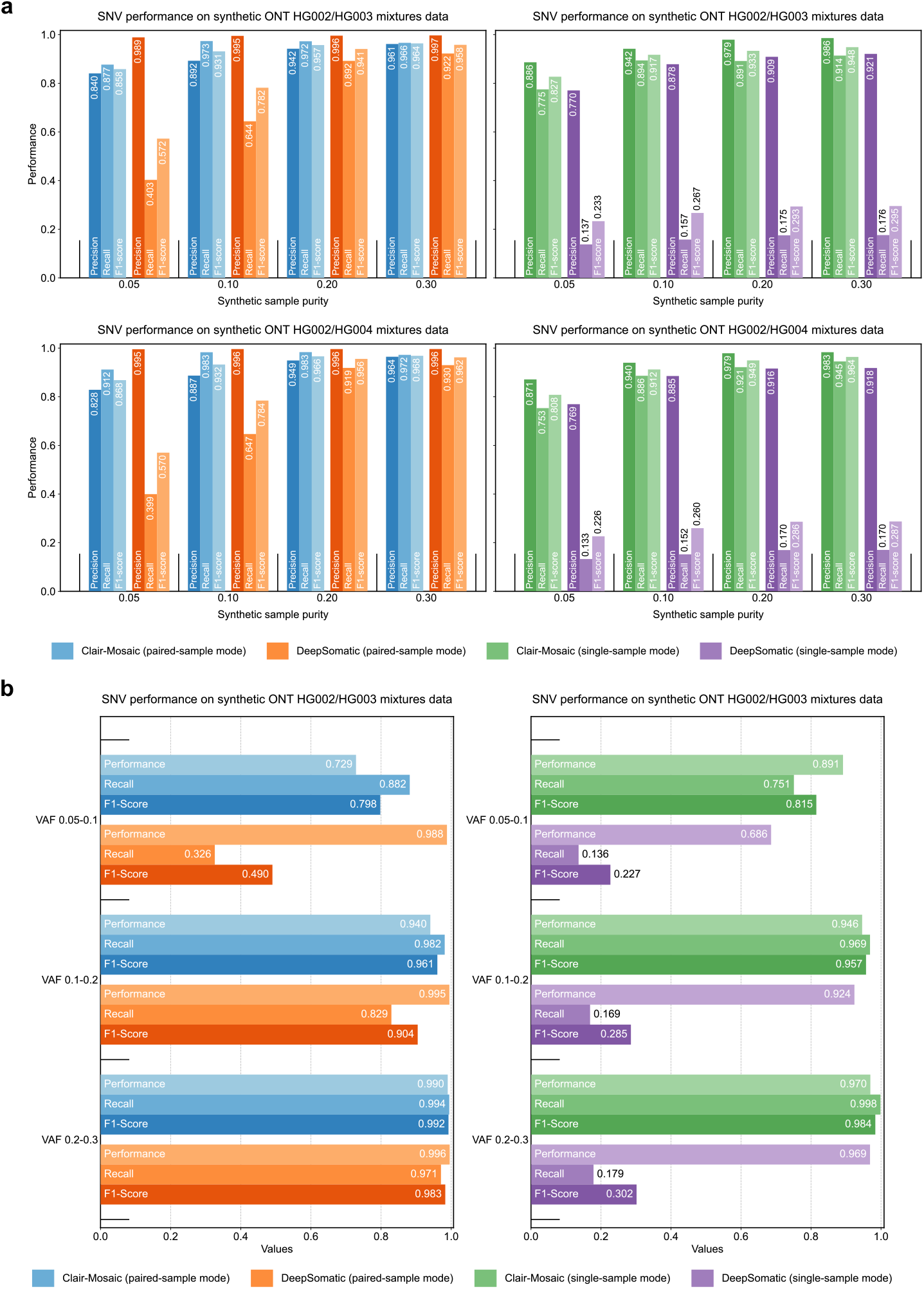
Performance in different sample purities and allelic fractions. **(a)** The figure illustrates the performance in different sample purities in HG002/HG003 and HG002/HG004 mixtures. **(b)** The figure illustrates the performance in different variant allelic fractions in HG002/HG003 synthetic datasets.

#### Performance at different VAF ranges

We further evaluated Clair-Mosaic across different VAF ranges using HG002/HG003 mixtures at varying sample purities (5%, 10%, 20%, 30%), with integrated results shown in **Figure 3b** and **Supplementary Table 3**. Performance was stratified into three VAF intervals: 5–10%, 10–20%, and 20–30%, to mimic different stages of mosaicism. In paired-sample mode, Clair-Mosaic achieved F1-scores of 79.78%, 96.05%, and 99.22% across the three VAF ranges, outperforming DeepSomatic by 30.77%, 5.60%, and 0.91%, respectively. In single-sample mode, Clair-Mosaic attained F1-scores of 81.52%, 95.72%, and 98.37% at the three VAF intervals. The performance gap was most pronounced in the low-VAF range (5–10%), underscoring Clair-Mosaic’s enhanced sensitivity in detecting mosaic variants with low allelic fractions. Furthermore, the method’s performance improved consistently with increasing VAF, highlighting the intrinsic challenge of detecting low-frequency variants.

#### Performance across various genomic contexts

To evaluate mosaic detection performance across various genomic contexts, we assessed Clair-Mosaic using challenging genomic regions defined by GIAB Stratifications v3.3^13^. We benchmarked the method on HG002/HG003 mixtures at 5%, 10%, 20%, and 30% sample purities. Regions were categorized into five classes according to GIAB stratification definitions: “Low complexity”, “Segmental duplications”, “Low mappability”, “Functional regions”, and “Other difficult regions”. As shown in **Figure 4a** and **Supplementary Table 4**, Clair-Mosaic consistently outperformed DeepSomatic in both paired-sample and single-sample modes across all genomic contexts. Despite this advantage, the absolute performance in challenging genomic contexts remained substantially below that of whole-genome benchmarks (86.93% versus 93.86% in paired-sample mode and 85.70% versus 91.93% in single-sample mode). These results highlight Clair-Mosaic’s capability in technically complex regions, while also underscoring the persistent challenge of accurately calling mosaic variants in complex genomic regions.

**Figure 4.**
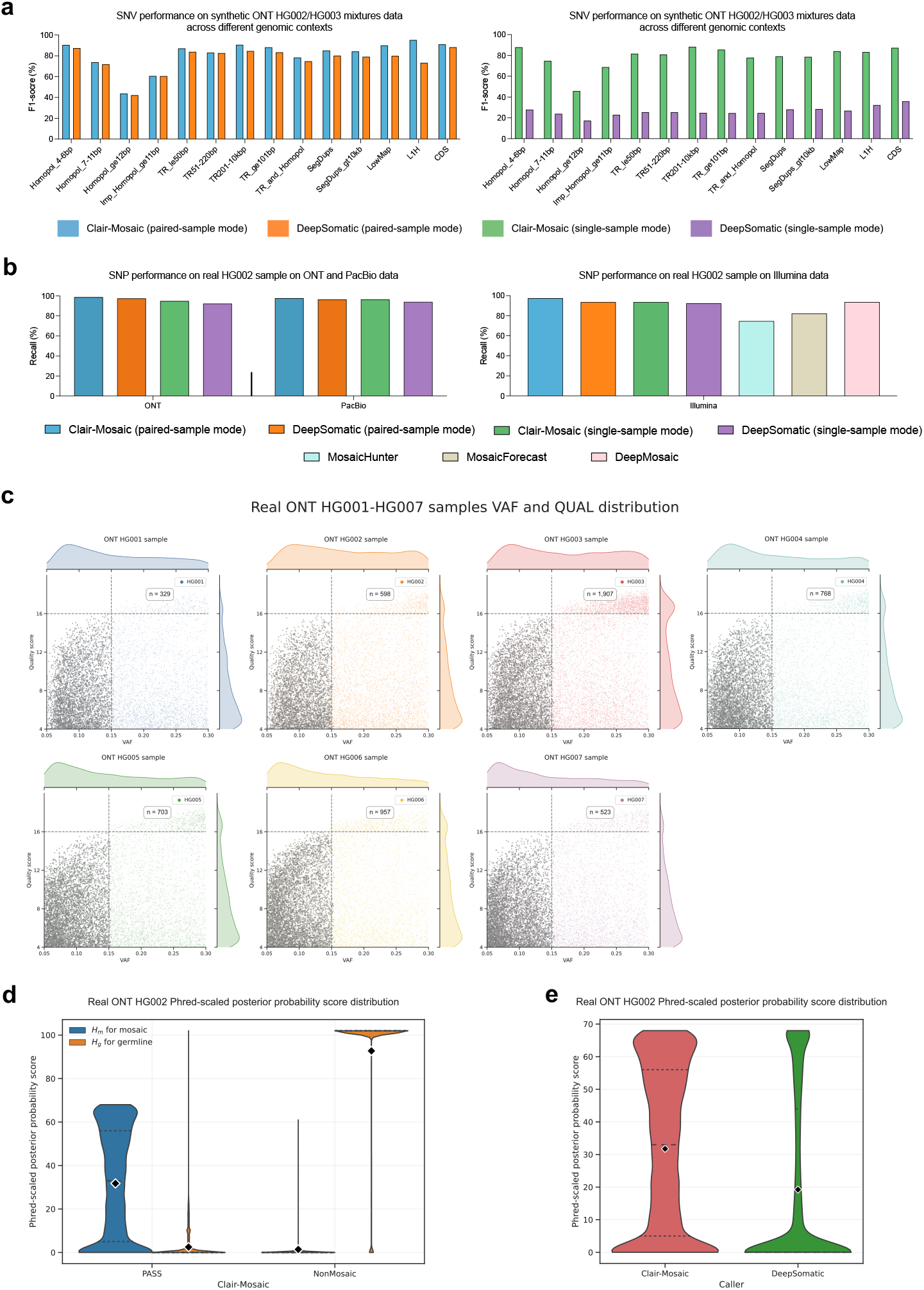
Performance on different genomic contexts and platforms. **(a)** The figure illustrates the performance of Clair-Mosaic on the ONT synthetic dataset across different genomic contexts, as defined by GIAB stratifications. **(b)** The figure illustrates the performance of different mosaic variant callers on the real HG002 dataset. **(c)** The distribution of quality scores and variant allelic fractions for mosaic variants called by Clair-Mosaic across the seven GIAB samples. **(d)** The distribution of Phred-scaled posterior probability scores for mosaic variants (the “PASS” category) and GRs-tagged germline variants (the “NonMosaic” category) called by Clair-Mosaic on the real HG002 dataset. **(e)** The distribution of Phred-scaled posterior probability scores for mosaic variants (the “PASS” category) called by Clair-Mosaic and DeepSomatic on the real HG002 dataset.

#### Performance using the real GIAB HG002 sample

In addition to synthetic benchmarks, we evaluated Clair-Mosaic on the real GIAB HG002 sample using the GIAB Consortium’s draft mosaic variant benchmark v1.0^25^, which provides 85 high-confidence mosaic SNVs orthogonally validated by multiple sequencing technologies. Among 85 SNVs, 7 variants with VAF below 5% were excluded from the final analysis. Given the limited number of benchmarkable mosaic variants, we focused on sensitivity as the primary metric to assess the capacity to detect true mosaic events. As shown in **Figure 4b** and **Supplementary Table 5**, Clair-Mosaic achieved a recall of 98.72% (77/78) in paired-sample mode and 94.87% (74/78) in single-sample mode. In contrast, DeepSomatic attained recalls of 97.44% (76/78) and 92.31% (72/78), respectively. The results demonstrate Clair-Mosaic’s capability for real-world mosaic variant detection and highlight its potential for practical clinical applications.

#### Performance on seven real GIAB samples (HG001–HG007)

To complement the analysis on HG002, we applied Clair-Mosaic to all seven GIAB samples (HG001–HG007) to investigate the distribution and consistency of detected mosaic variants across a diverse panel. As shown in **Figure 4c**, we analyzed the distribution of VAFs and quality scores for all candidate mosaic variants within the typical 5–30% VAF range. Using a quality cutoff (QUAL ≥16), we identified 329, 598, 1,907, 768, 703, 957, and 523 high-confidence mosaic variants in the ONT HG001–HG007 samples within the 15–30% VAF range, respectively. Although the absolute number of mosaic variants called in each sample varied, the counts were consistent with the expected range reported for normal samples.

#### Analysis of haplotype consistency checking on mosaic variant phasing

Leveraging the biological principle that postzygotic mosaic variants are confined to a single parental haplotype within affected cellular lineages^1^, Clair-Mosaic implements a haplotype-aware filtering strategy to reduce false positives by excluding candidate variants with allele support in both haplotypes. When applied to single-sample mode output from the real ONT HG002 sample, this approach tagged 15,248 variant candidates as sequencing artifacts, with two false positives shown in **Figure 5c**. Further validation against the GIAB HG002 draft mosaic benchmark (comprising 85 high-confidence SNVs) confirmed the biological relevance of this strategy. Of the called mosaic variants, 70 exhibited unambiguous haplotype consistency, being clearly assigned to a single phased haplotype, with two examples shown in **Figure 5b**. Eight variants could not be confidently phased due to insufficient phased reads in their genomic regions, while the remaining seven were excluded from haplotype analysis due to low read coverage or VAF below 0.05. These results demonstrate that haplotype consistency provides an effective strategy for mosaic variant authentication.

**Figure 5.**
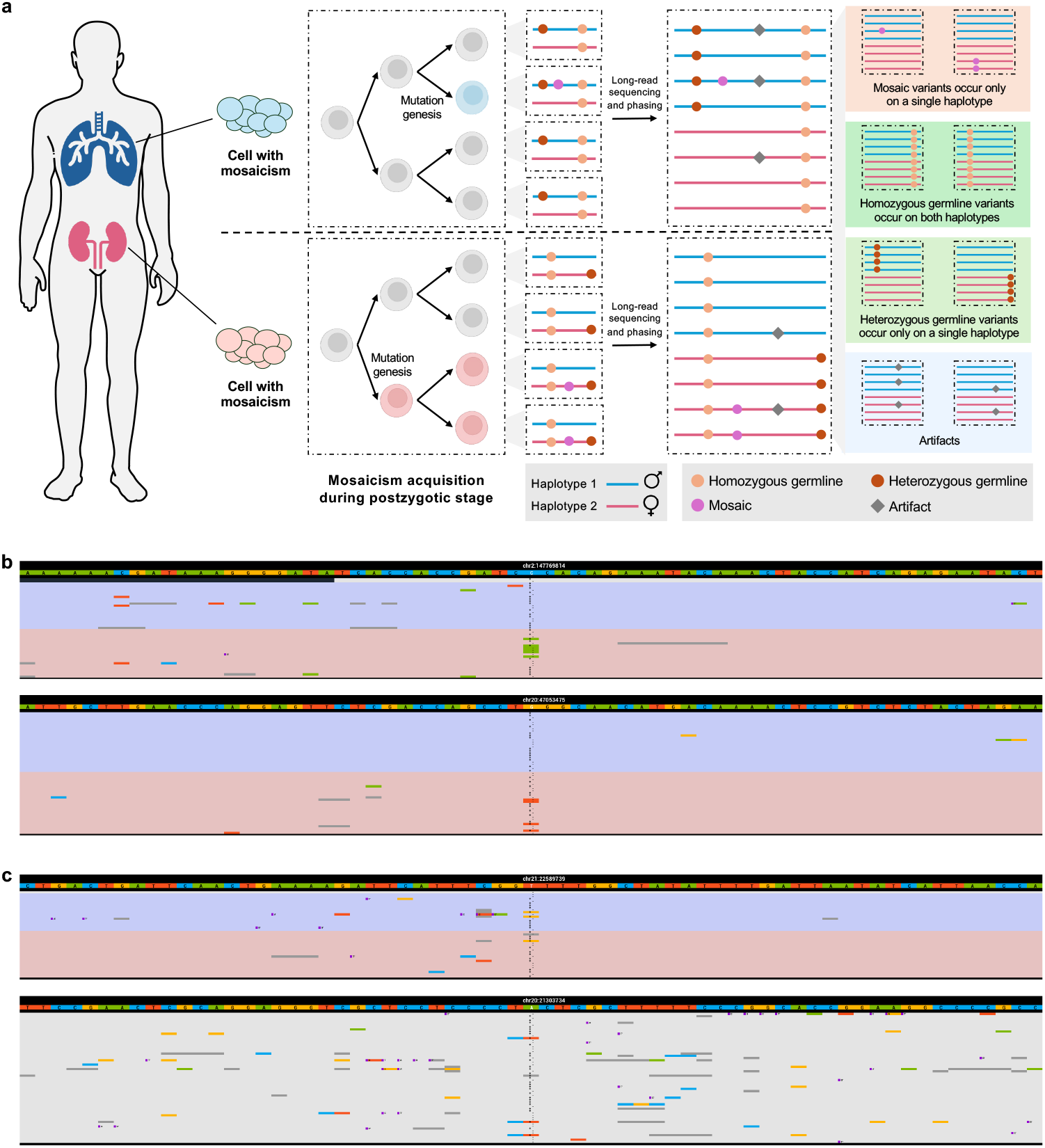
Haplotype consistency analysis on long reads. **(a)** The figure illustrates the biological basis for distinguishing true mosaic variants from sequencing artifacts using haplotype phasing. A true mosaic variant originates in a single somatic cell and propagates clonally through cell division, resulting in a haplotype-specific distribution. In contrast, sequencing artifacts appear randomly across haplotypes. **(b)** The figure illustrates the examples of a true mosaic variant existing in one haplotype, which is consistent with its occurrence. **(c)** The figure illustrates the examples of an artifact existing randomly in phased alignment or unphased alignment. The bases A, C, G, and T are depicted in green, blue, yellow, and red, respectively. The background in gray, purple, and pink represents an unknown haplotype, haplotype 1, and haplotype 2, respectively.

#### Analysis of probabilistic mosaic-germline discrimination using BayMGD

While GRs-based tagging effectively identifies common germline variants, it lacks the granularity to distinguish mosaic variants with intermediate allele frequencies (5–30%) from true germline polymorphisms. To address this limitation, Clair-Mosaic implements the Bayesian Mosaic-Germline Discriminator (BayMGD), a probabilistic framework that integrates observed individual allelic fractions, population allele frequency, and quality metrics to model continuous Phred-scaled probability scores for competing mosaic (*H*_*m*_) and germline (*H*_*g*_) hypotheses. Further details are provided in the **Methods – Bayesian Mosaic-Germline Discriminator for probabilistic variant classification** section.

As shown in **Figure 4d**, When applied to Clair-Mosaic’s single-sample mode results on the real ONT HG002 sample, BayMGD demonstrated high discriminatory consistency: called mosaic variants (the “PASS” category) showed higher Phred-scaled posterior probability scores for *H*_*m*_ (average: 31.75) compared with *H*_*g*_ (average: 2.48), whereas GRs-tagged germline variants (the “NonMosaic” category) exhibited higher probability scores for *H*_*g*_ (average: 92.71) than *H*_*m*_ (average: 1.40). We carried out the same analysis on DeepSomatic outputs, with results shown in **Figure 4e**, which yielded similar trends but with significantly lower *H*_*m*_ probability scores (average: 19.24), underscoring the effectiveness of the proposed BayMGD module. By applying quality and posterior probability scores thresholds (QUAL ≥16, *H*_*m*_ score ≥20), Clair-Mosaic tagged 57,293 low-confidence mosaic calls over 57,896 called mosaic variants in HG002. These results demonstrate BayMGD’s capability in identifying false positives caused by low-VAF germline variants.

#### Analysis of non-mosaic tagging using multiple germline resources

Clair-Mosaic employs four germline resources (GRs)—namely, gnomAD^30^, dbSNP^31^, the 1000 Genomes Project database^32^, and CoLoRSdb^33^—to systematically tag non-mosaic variants. We used a population allele frequency threshold of ≥0.001 to identify common germline variants, with more details described in the **Methods – Germline polymorphism tagging using multiple germline resources** section. When applied to Clair-Mosaic’s single-sample mode output for the real ONT HG002 sample, this multi-GR strategy tagged a total of 4,128,952 variants as germline-derived. Individual resources contributed 3,710,629 (gnomAD), 3,809,829 (dbSNP), 1,502,439 (1000 Genomes), and 4,084,056 (CoLoRSdb) tagged variants, with 1,443,475 germline variants identified concurrently by all four resources. These results demonstrate that the multi-GR approach is crucial for identifying the majority of germline polymorphisms, thereby significantly reducing false positives.

#### Analysis of mosaic variant tagging using the MosaicBase database

Clair-Mosaic further enhances variant interpretation by integrating the MosaicBase database^34^, a comprehensive resource of postzygotic mosaic variants curated from non-cancer disease cohorts and 422 healthy individuals, with more details described in the **Methods – Mosaic variant annotation using the MosaicBase database** section. Clair-Mosaic leverages MosaicBase to tag potential mosaic variants via exact allele matching and annotate corresponding gene and potential pathogenicity, thereby providing clinically enriched variant annotations. When applied to the real ONT HG002 output, 48 variants were annotated as existing in MosaicBase. For instance, the variant chr1:85,072,503 A->G was tagged as present in MosaicBase, associated with WDR63 Gene and annotated with potential relevance to Cockayne syndrome pathways. This approach bridges variant detection with biological interpretation, offering additional context to the reliability of the called mosaic variants.

### Performance on PacBio data

To validate the generalizability of Clair-Mosaic, we evaluated its performance on the PacBio platform, with a summary of datasets provided in **Supplementary Table 1**. The PacBio models were trained on synthetic datasets generated from biologically independent HG004 and HG003 samples sequenced on the PacBio Revio system. We then performed benchmarking on synthetic HG002/HG003 and HG002/HG004 mixtures at different sample purities, where 50-fold was used for input and 25-fold for control. As shown in **Figure 6a** and **Supplementary Table 5**, Clair-Mosaic achieved F1-scores of 98.11%, 98.97%, 98.93%, and 98.82% at 5%, 10%, 20%, and 30% purity levels in paired-sample mode for HG002/HG003 mixtures, outperforming DeepSomatic by 6.04%, 3.03%, 1.12%, and 0.43%, respectively. Similar trends were observed for HG002/HG004 mixtures. In single-sample mode, Clair-Mosaic achieved F1-scores of 95.20%, 97.56%, 97.89%, and 98.36% at the four purity levels. Additionally, we evaluated performance on the real PacBio HG002 50-fold dataset sequenced using the PacBio Revio system. As shown in **Figure 4b** and **Supplementary Table 7**, Clair-Mosaic achieved a recall of 97.62% in paired-sample mode and 96.43% in single-sample mode, consistently surpassing DeepSomatic. These multi-platform validation results demonstrate Clair-Mosaic’s capability in detecting mosaicism independent of sequencing technology, establishing it as a versatile and reliable tool for long-read mosaic variant detection.

**Figure 6.**
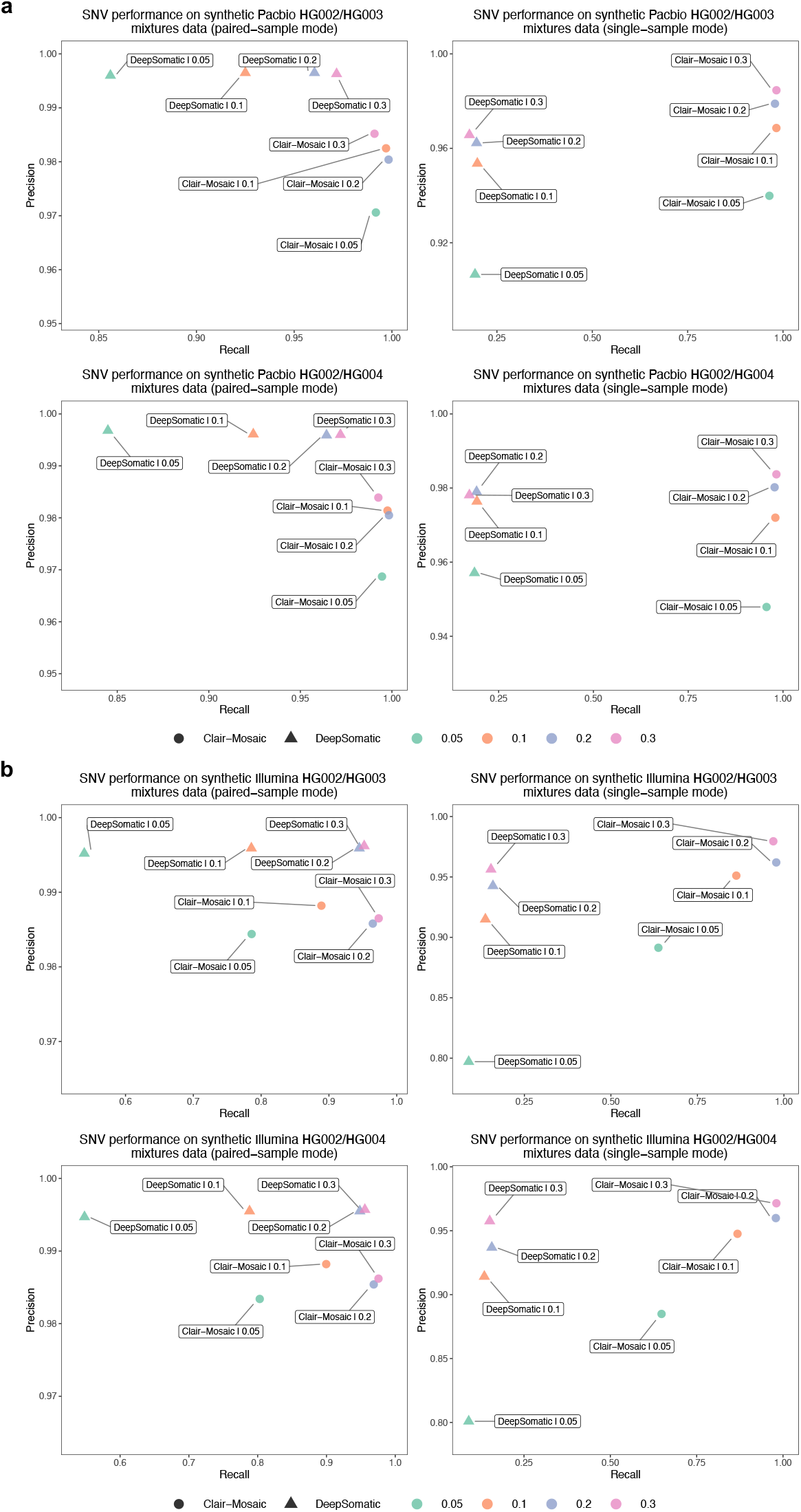
Performance analysis on the PacBio and Illumina platforms. **(a)** The precision and recall performance of different callers on the PacBio dataset. **(b)** The precision and recall performance of different callers on the Illumina dataset. Different colors of the dots represent different sample purities.

### Performance on Illumina data

Short-read mosaic variant calling has been extensively studied, leading to the development of numerous notable methods such as MosaicHunter^21^, MosaicForecast^22^, and DeepMosaic^23^. Although Clair-Mosaic was primarily designed for long-read sequencing data, its framework is not limited to long reads. Except for haplotype consistency checking, which is disabled due to insufficient read length in short reads, Clair-Mosaic’s core methodology remains applicable to short-read data. To test this, for the Illumina platform, we trained the model on synthetically mixed data generated from biologically independent HG004 and HG003 samples sequenced on Illumina’s HiSeq X and NovaSeq 6000 platforms; more details are provided in **Supplementary Table 1**. We then performed benchmarking on synthetic HG002/HG003 and HG002/HG004 mixtures across varying sample purities. As illustrated in **Figure 6b** and **Supplementary Table 6**, Clair-Mosaic achieved F1-scores of 87.41%, 93.59%, 97.52%, and 97.99% at 5%, 10%, 20%, and 30% purity levels in paired-sample mode for HG002/HG003 mixtures, exceeding DeepSomatic by an average of 6.10% across all purity levels. Evaluations in single-sample mode showed similar advantages, with F1-scores of 74.32%, 90.46%, 96.98%, and 97.44% at the four purity levels. Similar trends were observed for HG002/HG004 mixtures. Evaluation on the real Illumina HG002 sample further confirmed Clair-Mosaic’s applicability to short-read data, with Clair-Mosaic achieving a recall of 93.67% in single-sample mode, consistently exceeding or meeting the performance of current state-of-the-art short-read mosaic callers, including MosaicHunter (74.68%), MosaicForecast (82.28%), DeepMosaic (93.67%), and DeepSomatic (92.41%). These results demonstrate that Clair-Mosaic provides a unified framework for mosaic variant detection, delivering robust performance across sequencing technologies that is comparable to specialized methods.

### Indel performance analysis

Compared to SNV detection, mosaic Indel detection presents distinct challenges, primarily due to elevated error rates in complex genomic regions (e.g., homopolymers and tandem repeats) inherent to long-read technologies^13^, which generate substantial sequencing artifacts that obscure true low-VAF mosaic Indels. Furthermore, the extreme scarcity of mosaic Indels compared to SNVs severely hinders their evaluation. In light of these limitations, we evaluated mosaic Indel calling performance using only synthetic datasets. Specifically, we benchmarked Clair-Mosaic on HG002/HG003, HG002/HG004, HG005/HG006, and HG005/HG007 synthetic mixtures across varying sample purities. As shown in **Figure 7, Supplementary Figure 2**, and **Supplementary Table 2**, Clair-Mosaic achieved F1-scores of 32.93%, 54.95%, 70.76%, and 75.41% at 5%, 10%, 20%, and 30% purity levels in paired-sample mode for HG002/HG003 mixtures, outperforming DeepSomatic by an average of 20.12%. Similar trends were observed for HG002/HG004, HG005/HG006, and HG005/HG007 mixtures. In single-sample mode, Clair-Mosaic attained F1-scores of 29.99%, 49.75%, 65.54%, and 71.39% at the purity levels. Although these results demonstrate Clair-Mosaic’s relative advantage in Indel calling, its absolute performance remains constrained, with a maximum F1-score of 75.82%, which is significantly lower than that for SNVs. This underscores that the reliable detection of mosaic Indels remains a significant challenge.

**Figure 7.**
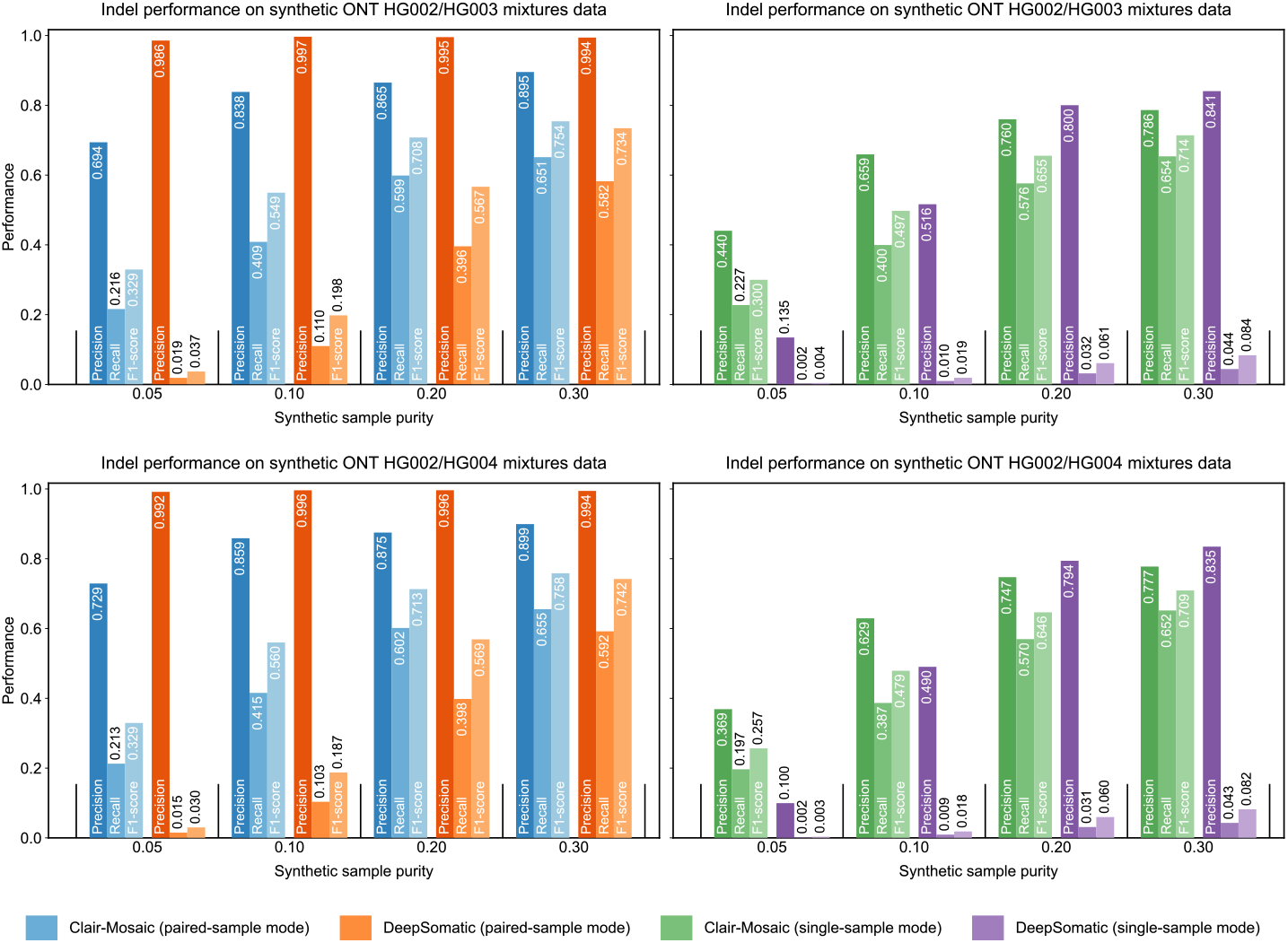
Indel performance analysis. The figure illustrates the Indel performance of different callers in different synthetic mixtures.

### Analysis of mosaic variants called uniquely in long read

Given the demonstrated superiority of long-read technologies for providing ambiguous alignment in complex genomic regions^26^, we sought to investigate the specific mosaic variants uniquely identified by them. All mosaic variants called by ONT and PacBio that were absent from the Illumina data were collected and categorized according to GIAB genomic stratifications by various genomic contexts. As shown in **Figure 8**, ONT and PacBio detected 3,096, 3,166, and 2,491 mosaic variants missed by short-read sequencing in GIAB samples HG002, HG003, and HG004, respectively. Within the difficult genomic regions defined by GIAB, tandem repeats/homopolymers, segmental duplications, and low-mappability regions accounted for 72.5%, 66.8%, and 62.3% of these variants, respectively. This result underscores the enhanced capability of long-read sequencing for identifying mosaic variants in challenging genomic contexts that are typically missed by conventional short-read sequencing.

**Figure 8.**
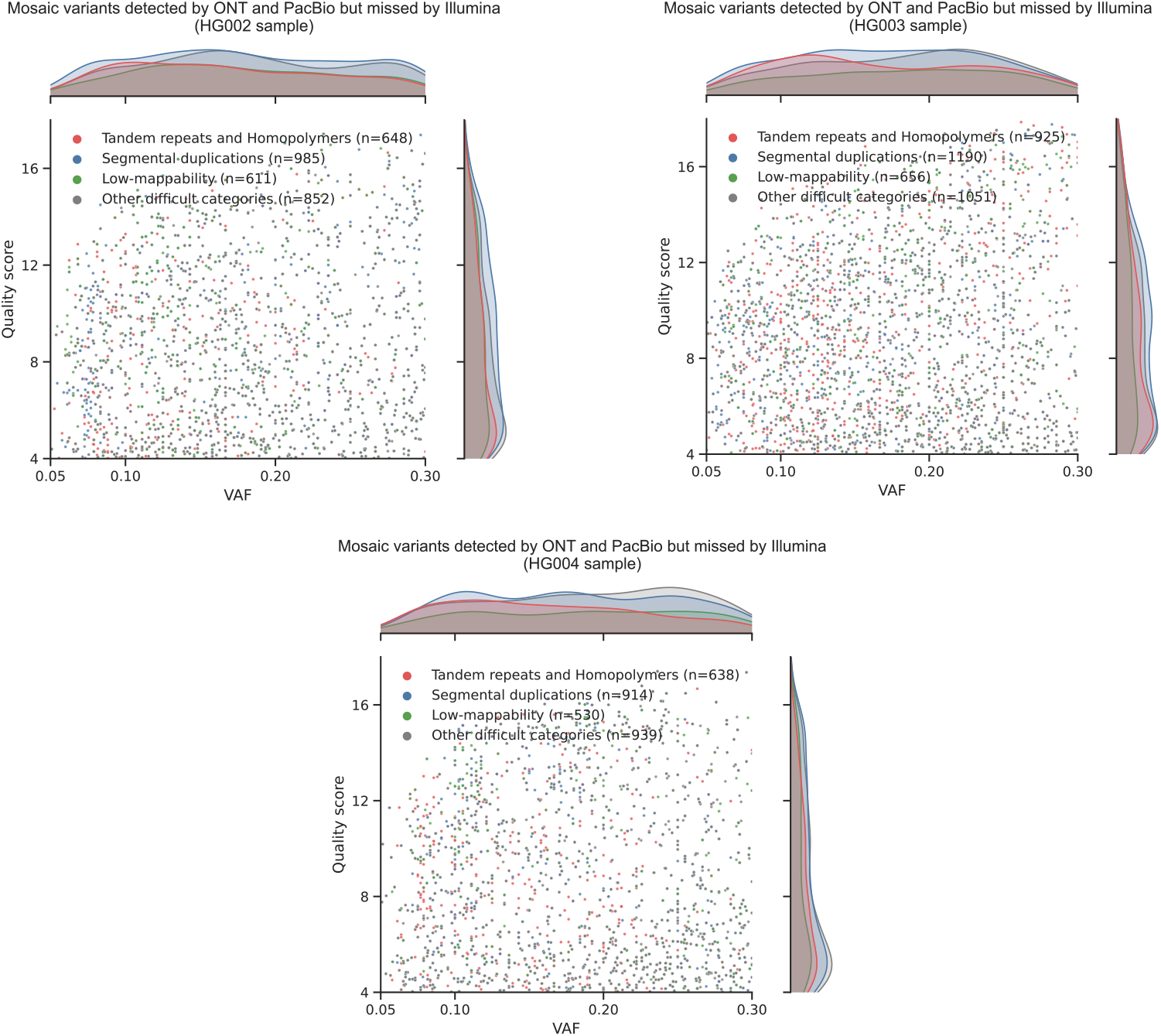
Analysis of mosaic variants called by long read but missed in short read. The quality score and VAF distribution of the mosaic variants called by ONT and PacBio but missed in Illumina stratified by different GIAB genomic contexts on 50-fold ONT HG002, HG003, and HG004 datasets.

## Discussion

Somatic mosaicism has emerged as a crucial area of genomic research, with broad implications for understanding health, aging, and disease mechanisms. The recent launch of the Somatic Mosaicism Across Human Tissues (SMaHT) Network^2^ has marked a major advance, aiming to catalog somatic mutations across dozens of tissue types from hundreds of non-diseased donors. This initiative will systematically map the human body’s mutational landscape, illuminating how mutations accumulate and contribute to health and disease. Its scale underscores the urgent need for developing robust mosaic detection methods, particularly those leveraging long-read sequencing.

To address this need, we present Clair-Mosaic, a deep-learning-based method for detecting mosaic small variants from long-read sequencing data. Clair-Mosaic supports both paired-sample and single-sample analyses, enhancing its utility across diverse research and clinical contexts where matched controls may be unavailable. A key innovation is its training on hundreds of millions of synthetic variants spanning a wide spectrum of read coverages, allelic fractions, and genomic contexts. The method leverages the long-range information of sequencing reads, which span tens of kilobases, to perform haplotype-aware filtering that effectively excludes sequencing artifacts. Furthermore, Clair-Mosaic incorporates a Bayesian Mosaic-Germline Discriminator, which uses population allele frequency priors to distinguish sample-specific mosaic variants from common germline polymorphisms. Together, these features robustly eliminate major germline variants and technical artifacts, enabling highly accurate mosaic variant detection. We expect that the development of reliable mosaic variant callers will substantially improve our understanding of how somatic mosaicism influences human health, aging, and disease.

Despite these advances, our study underscores two key challenges. First, the field still lacks sufficient real mosaic variant datasets for benchmarking with long-read sequencing. This critical gap is exacerbated by the intrinsic biological scarcity of mosaic variants within any individual genome, which fundamentally limits the scale and diversity of variants available for method evaluation. Although synthetic data enable scalable training of deep learning models, such *in silico* mixtures cannot fully replicate the biological complexity of real mosaic variants, including tissue-specific noise, structural variations, and mosaic-specific error profiles. We anticipate that the release of Clair-Mosaic will facilitate the detection of mosaic variants across diverse samples, thereby advancing our understanding of somatic mosaicism in human biology. Second, accurate mosaic Indel detection remains challenging. Due to higher error rates and alignment uncertainties in complex genomic regions, current Indel calling performance is less robust than that for SNVs, limiting its practical utility.

## Methods

### Overview of Clair-Mosaic

As shown in **Figure 1a**, Clair-Mosaic is designed to detect the postzygotic mosaic variants sequenced in long reads. **Figure 1b** provides an overview of the Clair-Mosaic variant calling workflow. Clair-Mosaic supports two calling modes: a paired-sample mode and a single-sample mode. In paired-sample mode, both an input sample and a control sample are provided with read alignments in BAM/CRAM format for identifying candidate mosaic variants supported by sufficient read evidence and allele alterations. The variant candidates and their corresponding flanking window sites are summarized into pileups, which are then concatenated vertically with the aligned positions from the control sample to create tensors. Using the pileup model for inference, Clair-Mosaic outputs three probabilities indicating whether a variant candidate is a mosaic variant, a germline variant, or a sequencing artifact. In real-world scenarios, a matched control sample is often unavailable. Therefore, Clair-Mosaic also supports a single-sample mode that processes only the input sample. Similarly, variant candidates from the input sample are extracted using a heuristic strategy based on alternate allele read support, summarized into pileups, and converted into tensors. Clair-Mosaic then outputs two probabilities representing the likelihood of a variant candidate being a mosaic variant or an artifact. Following network prediction, several post-processing steps—including haplotype consistency checking, filtering against germline resources, and the Bayesian Mosaic-Germline Discriminator (BayMGD)—are applied to filter out non-mosaic variants and eliminate sequencing artifacts, yielding more reliable mosaic calling results. Additionally, Clair-Mosaic includes a mosaic-specific tagging module that annotates known mosaic variants with corresponding gene and potential pathogenicity information from the MosaicBase database.

### Clair-Mosaic’s data synthesis strategy

To overcome the scarcity of real mosaic variants for model training and performance benchmarking, Clair-Mosaic employs a synthetic data generation strategy that combines sequencing reads from distinct samples by treating germline variants unique to one sample as mosaic variants in synthetic mixtures. This approach ensures that the synthetic data captures the full spectrum of VAF distributions, particularly the predominance of low-VAF variants characteristic of real mosaicism. Critically, while the mosaic variants are synthetic, the sequencing artifacts reflect real error profiles, ensuring that a model trained on synthetic data performs well on real data.

For synthetic benchmarking, Clair-Mosaic leverages biologically related family trio samples (HG002/HG003/HG004 and HG005/HG006/HG007), aligning with the principle that postzygotic mosaic variants are not be inherited. We used the child as the input sample and one of the parents as the control sample for data synthesis, resulting in four input/control pairs: HG002/HG003, HG002/HG004, HG005/HG006, and HG005/HG007 for evaluation. Synthetic mixtures were set at 5%, 10%, 20%, and 30% purity levels to simulate biologically relevant VAF ranges (5–30%), with truth sets constructed by subtracting the control sample’s germline variants from those of the input sample.

For model training, Clair-Mosaic uses biologically independent GIAB samples (e.g., HG002 and HG001) to enhance generalizability and robustness. Specifically, using GIAB HG002 as the input mosaic sample source and GIAB HG001 as the matched control sample source, the alignments of both samples were split into smaller non-overlapping chunks of ~4-fold coverage each. These chunks were then systematically recombined to simulate different sequencing coverages (by controlling the total number of chunks) and different VAFs (by adjusting the proportion of chunks from each sample). During synthesis, three variant categories were defined for model training: “Mosaic” (germline variants present in HG002 but absent in HG001), “Germline” (germline variants shared by HG002 and HG001), and “Artifact” (candidate variants in the synthetic mixture not supported by the truth germline variants of HG002). This strategy produces datasets containing millions of variant candidates, providing the volume and diversity necessary to train a deep neural network.

### Clair-Mosaic’s input and output

#### Input

Clair-Mosaic takes read alignments—from either a paired input and control sample (in paired-sample mode) or a single input sample (in single-sample mode)—along with the corresponding reference genome as input. The input read alignments are processed in parallel to accelerate tensor generation, model inference, and post-processing, resulting in a final VCF output. Clair-Mosaic’s pileup input comprises 1,122 integers, covering 33 genomic positions (the candidate variant position plus 16 flanking positions on each side) with 34 features per position. The 34 pileup features provide statistics information to summarize both the forward (+) and reverse (-) strand of sequencing reads of 1) nucleotides (A+/C+/G+/T+ and A-/C-/G-/T-), 2) insertions (I_S_+/I_1S_+ and I_S_-/I_1S_-), 3) deletions ((D_S_+/D_1S_+/D_R_+ and D_S_-/D_1S_-/D_R_-)), 4) nucleotides with low mapping quality (A_LMQ_+/C_LMQ_+/G_LMQ_+/T_LMQ_+ and A_LMQ_-/C_LMQ_-/G_LMQ_-/T_LMQ_-), and 5) nucleotides with low base quality (A_LBQ_+/C_LBQ_+/G_LBQ_+/T_LBQ_+ and A_LBQ_-/C_LBQ_-/G_LBQ_-/T_LBQ_-), with detailed description described in the **Supplementary Information – Pileup input**.

#### Clair-Mosaic’s model architecture and output task

Clair-Mosaic uses a similar pileup model architecture for both paired-sample and single-sample inputs. In paired-sample mode, the model employs two bidirectional gated recurrent unit (Bi-GRU) layers to encode sequential pileup features generated by vertically concatenating the input and control sample features. The paired-sample model outputs three probabilities, indicating whether a variant candidate is a mosaic variant, a germline variant, or a sequencing artifact. Notably, with the control sample as a reference, Clair-Mosaic can distinguish mosaic variants from germline variants based on their presence: mosaic variants are typically absent in the control, whereas germline variants are present in both the input and control samples. In single-sample mode, Clair-Mosaic uses a convolutional vision transformer (CvT)^35^-based pileup model to estimate the likelihood of a candidate being a mosaic variant versus a sequencing artifact. Based on these probabilities, Clair-Mosaic then classifies each candidate as either a mosaic variant or an artifact.

#### VCF output

Clair-Mosaic outputs results in VCF format, with variants categorized into three types: 1) variants labeled as “PASS” that pass all filters and quality score thresholds; 2) variants labeled as “LowQual” due to low variant quality scores (QUAL < 4 by default, configurable via command-line options) or those filtered out by post-processing filters; and 3) variants tagged as “NonMosaic” if they are present in any of the specified germline resources. For each variant, supporting information such as allelic fraction, read depth, and allele depth is included in the VCF file. In paired-sample calling mode, the corresponding read support information from the control sample is also provided.

### Haplotype consistency checking for low-VAF mosaic variants

Clair-Mosaic leverages the inherent haplotype-resolving capability of long-read sequencing to distinguish true mosaic variants from sequencing artifacts. The rationale is that mosaic variants typically arise postzygotically and are therefore confined to a single parental haplotype (either paternal or maternal) within affected cellular lineages^1^, as illustrated in **Figure 5a**. Based on the premise that true mosaics are haplotype-restricted, Clair-Mosaic distinguishes them from artifacts, which show no haplotype specificity.

The workflow begins by applying Clair3^16^ to detect heterozygous germline SNP variants. These germline variants are phased using LongPhase^36^, and the phased variants are then used to assign reads to specific haplotypes (read haplotagging). For each variant candidate, Clair-Mosaic calculates the alternative allele VAF separately within reads assigned to each haplotype. Variants failing strict phasing criteria—such as those exhibiting significant alternative allele support in both haplotypes (defined as having >2 alternative allele-supporting reads or an alternative VAF >0.05 in the minor haplotype)—are filtered out as false positives. Moreover, variants located in poorly phased regions with insufficient haplotagged reads are excluded from analysis due to inconclusive phasing evidence. This haplotype-based filtering enables high-specificity variant detection, especially for low-VAF mosaic variants.

### Germline polymorphism tagging using multiple germline resources

Distinguishing true mosaic variants from germline variants is challenging, especially in the absence of a matched control sample. Following a common practice in somatic variant calling^27, 37, 38^, Clair-Mosaic supports the use of germline resources (GRs) to tag non-mosaic variants. By default, Clair-Mosaic integrates four germline variant databases: gnomAD^30^, dbSNP^31^, the 1000 Genomes Project (1000G)^32^, and the Consortium for Long-Read Sequencing (CoLoRSdb) database^33^. The original gnomAD r2.1, dbSNP v138, and 1000G databases were sourced from the GATK resource bundle, while the inclusion of CoLoRSdb was inspired by Park et al.^27^. After applying a population allele frequency threshold of VAF ≥ 0.001, a total of 35,551,905, 60,683,019, 2,609,566, and 49,550,902 variants from these respective databases were retained for constructing the GRs used by Clair-Mosaic to tag non-mosaic variants.

### Bayesian Mosaic-Germline Discriminator for probabilistic variant classification

While GRs-based tagging effectively filters common germline polymorphisms, it operates using manually defined allele frequency thresholds and lacks the capacity to distinguish mosaic variants with intermediate allele frequencies (e.g., 5%–30%) from rare germline polymorphisms. To address this issue, we developed a **Bay**esian **M**osaic-**G**ermline **D**iscriminator (BayMGD), a probabilistic framework that models multiple evidence sources to compute continuous probability scores for mosaic and germline hypotheses (denoted as *H*_*m*_ and *H*_*g*_, respectively). The posterior probability for each hypothesis given evidence *E* is formulated as:

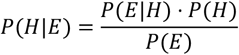

where the likelihood *P*(*E*|*H*) integrates three evidence sources (observed individual allelic fraction (*AF*_*obs*_), population frequency from gnomAD (*AF*_*pop*_), and quality score (*QUAL*) under conditional independence:

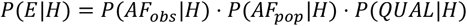

Prior probabilities reflect the biological prevalence of each variant category, with *P*(*H*_*m*_) = 0.05 and *P*(*H*_*g*_) = 0.95. Likelihood functions were tailored to each evidence type: *AF*_*obs*_ for mosaic hypothesis *H*_*m*_ was modeled using a Beta distribution (α=3, β=30; peak ~9%) to capture low-VAF signatures, while *AF*_*obs*_ for germline hypothesis *H*_*g*_ followed a normal distribution (μ=0.5, σ=0.18) to accommodate natural variation with averaged 50% heterozygosity. The likelihoods of *AF*_*pop*_ used sparse Beta distributions (for *H*_*m*_: α=1.2, β=200; for *H*_*g*_: α=2, β=2) to emphasize the rarity of mosaic variants in population databases. *QUAL* was modeled via log-normal distributions (μ=3.0, σ=1.5) for both hypotheses. The parameters for all distributions were manually selected. Posterior probabilities were transformed into Phred-scaled scores for clinical interpretation. Based on the computed probability scores, a candidate variant is tagged as germline if the *H*_*m*_ probability score exceeds that of *H*_*g*_.

### Mosaic variant annotation using the MosaicBase database

MosaicBase^34^ is a comprehensive database of postzygotic mosaic variants curated from non-cancer disease cohorts and healthy individuals. It comprises 6,698 mosaic variants associated with 266 non-cancer diseases and 27,991 mosaic variants identified in 422 healthy individuals, with detailed genomic and phenotypic information annotated for each entry. Clair-Mosaic utilizes MosaicBase to annotate called variants via exact allele matching. Variants present in the database are tagged with corresponding gene and potential pathogenicity information, thereby facilitating biological and clinical interpretation.

### Benchmarking metrics

We benchmarked Clair-Mosaic on both synthetic GIAB datasets and the real GIAB HG002 sample. For evaluations on synthetic data, we generated *in silico* mixed samples from two independent family trios: HG002/HG003/HG004 Ashkenazi trio and HG005/HG006/HG007 Chinese trio^13, 26^. For each trio, sequencing reads from the child sample (e.g., HG002 or HG005) were added into their parent sample (e.g., HG003 or HG006) at varying sample purities, treating the latter as the control. The truth variant sets were constructed by subtracting germline variants of the control sample from those of the input sample, thereby regarding sample-specific germline variants as synthetic mosaic events. The benchmarking regions were defined as the intersection of GIAB high-confidence regions between two mixed samples. For evaluation on real data, we utilized the HG002 draft mosaic variant benchmark v1.0 released by GIAB Consortium^25^, which provides a truth VCF containing 85 high-confidence mosaic SNVs. Performance was assessed using the Clair-Mosaic’s “compare_vcf” submodule to compute precision, recall, and F1-score for SNV and Indel. All commands and parameters used for executing the benchmarked callers are provided in the **Supplementary Information – Command lines used** section.

### Computational performance

Clair-Mosaic is implemented in Python and PyTorch, with computational components accelerated via PyPy. Model training was conducted on GPUs with VRAM ≥6 GB, facilitated by a lightweight pileup-based architecture. Total training time for the dual-mode architecture was completed in approximately 6 hours using an NVIDIA RTX 3090 GPU; compatibility was also verified on other GPUs, including the RTX 2080 Ti and 4090. For model inference, Clair-Mosaic requires only CPUs for computing. Peak memory usage remained below 1 GB per CPU process.

## Code availability

Clair-Mosaic is open-source and available at https://github.com/HKU-BAL/Clair-Mosaic under the BSD 3-Clause license. The evaluation results in this paper were based on the Clair-Mosaic v0.0.1 release. A public Docker image hkubal/clair-mosaic:latest is also available for simplicity to evaluate Clair-Mosaic’s pipeline.

## Data availability

All data used in this study, including the 1) ONT, PacBio, and Illumina sequencing datasets; 2) GIAB HG002 truth mosaic variant set; 2) GIAB HG001-HG007 truth germline variant sets; and 3) reference genomes, are available in the **Supplementary Information – Data availability** section. All analysis output, including the VCFs, are available at http://www.bio8.cs.hku.hk/clair-mosaic/analysis_result.

## Acknowledgements

R.L. was supported by CRF (C7003-24Y) of the Research Grants Council (RGC) of Hong Kong, and the URC fund at HKU.

## Author contributions

R. L. conceived the study. L. C., Z. Z., and R. L. designed the algorithms, implemented and benchmarked Clair-Mosaic, and wrote the paper. M. H., A. O. K. W., and X. Y. evaluated the benchmarking results. All authors revised the manuscript.

## References

1. Freed, D., Stevens, E.L. & Pevsner, J. Somatic Mosaicism in the Human Genome. Genes 5, 1064–1094 (2014).

2. Coorens, T.H.H. et al. The Somatic Mosaicism across Human Tissues Network. Nature 643, 47–59 (2025).

3. Dou, Y.M., Gold, H.D., Luquette, L.J. & Park, P.J. Detecting Somatic Mutations in Normal Cells. Trends in Genetics 34, 545–557 (2018).

4. Biesecker, L.G. & Spinner, N.B. A genomic view of mosaicism and human disease. Nature Reviews Genetics 14, 307–320 (2013).

5. Poduri, A., Evrony, G.D., Cai, X.Y. & Walsh, C.A. Somatic Mutation, Genomic Variation, and Neurological Disease. Science 341, 1237758 (2013).

6. Breuss, M.W. et al. Somatic mosaicism reveals clonal distributions of neocortical development. Nature 604, 689–696 (2022).

7. Martincorena, I. et al. Somatic mutant clones colonize the human esophagus with age. Science 362, 911–917 (2018).

8. Vijg, J. & Dong, X. Pathogenic Mechanisms of Somatic Mutation and Genome Mosaicism in Aging. Cell 182, 12–23 (2020).

9. Lim, E.T. et al. Rates, distribution and implications of postzygotic mosaic mutations in autism spectrum disorder. Nature Neuroscience 20, 1217–1224 (2017).

10. Freed, D. & Pevsner, J. The Contribution of Mosaic Variants to Autism Spectrum Disorder. Plos Genetics 12 (2016).

11. Kim, J.H. et al. Analysis of low-level somatic mosaicism reveals stage and tissue-specific mutational features in human development. Plos Genetics 18 (2022).

12. Ameur, A., Kloosterman, W.P. & Hestand, M.S. Single-Molecule Sequencing: Towards Clinical Applications. Trends in Biotechnology 37, 72–85 (2019).

13. Wagner, J. et al. Benchmarking challenging small variants with linked and long reads. Cell Genom 2 (2022).

14. Logsdon, G.A., Vollger, M.R. & Eichler, E.E. Long-read human genome sequencing and its applications. Nature Reviews Genetics 21, 597–614 (2020).

15. Olson, N.D. et al. Variant calling and benchmarking in an era of complete human genome sequences. Nature Reviews Genetics 24, 464–483 (2023).

16. Zheng, Z.X. et al. Symphonizing pileup and full-alignment for deep learning-based long-read variant calling. Nature Computational Science 2, 797–803 (2022).

17. Poplin, R. et al. A universal SNP and small-indel variant caller using deep neural networks. Nature Biotechnology 36, 983–987 (2018).

18. Luo, R.B., Sedlazeck, F.J., Lam, T.W. & Schatz, M.C. A multi-task convolutional deep neural network for variant calling in single molecule sequencing. Nat Commun 10 (2019).

19. Luo, R.B. et al. Exploring the limit of using a deep neural network on pileup data for germline variant calling. Nat Mach Intell 2, 220–227 (2020).

20. Shafin, K. et al. Haplotype-aware variant calling with PEPPER-Margin-DeepVariant enables high accuracy in nanopore long-reads. Nat Methods 18, 1322–1332 (2021).

21. Huang, A.Y. et al. MosaicHunter: accurate detection of postzygotic single-nucleotide mosaicism through next-generation sequencing of unpaired, trio, and paired samples. Nucleic Acids Research 45 (2017).

22. Dou, Y.M. et al. Accurate detection of mosaic variants in sequencing data without matched controls. Nature Biotechnology 38, 314-+ (2020).

23. Yang, X.X. et al. Control-independent mosaic single nucleotide variant detection with DeepMosaic. Nature Biotechnology 41, 870–877 (2023).

24. Ha, Y.J. et al. Comprehensive benchmarking and guidelines of mosaic variant calling strategies. Nat Methods 20 (2023).

25. Daniels, C.A. et al. A robust benchmark for detecting low-frequency variants in the HG002 Genome In A Bottle NIST reference material. bioRxiv (2024).

26. Zook, J.M. et al. A robust benchmark for detection of germline large deletions and insertions. Nature Biotechnology 38, 1347–1355 (2020).

27. Park, J. et al. Accurate somatic small variant discovery for multiple sequencing technologies with DeepSomatic. Nature Biotechnology (2025).

28. Zhenxian Zheng, J.S., Lei Chen, Yan-Lam Lee, Tak-Wah Lam, Ruibang Luo ClairS: a deep-learning method for long-read somatic small variant calling. bioRxiv (2023).

29. Nanopore EPI2ME Labs, https://github.com/epi2me-labs/wf-somatic-variation. (2023).

30. Gudmundsson, S. et al. Variant interpretation using population databases: Lessons from gnomAD. Human Mutation 43, 1012–1030 (2022).

31. Sherry, S.T. et al. dbSNP: the NCBI database of genetic variation. Nucleic Acids Research 29, 308–311 (2001).

32. Siva, N. 1000 Genomes project. Nature Biotechnology 26, 256–256 (2008).

33. Lake, J.A. & Consortium of Long Read Sequencing (CoLoRS). Consortium of Long Read Sequencing Database (CoLoRSdb) (v1.1.0) [Data set]. Zenodo. (2024).

34. Yang, X.X. et al. MosaicBase: A Knowledgebase of Postzygotic Mosaic Variants in Noncancer Disease-related and Healthy Human Individuals. Genomics Proteomics & Bioinformatics 18, 140–149 (2020).

35. Wu, H.P. et al. CvT: Introducing Convolutions to Vision Transformers. 2021 Ieee/Cvf International Conference on Computer Vision (Iccv 2021), 22–31 (2021).

36. Lin, J.H., Chen, L.C., Yu, S.C. & Huang, Y.T. LongPhase: an ultra-fast chromosome-scale phasing algorithm for small and large variants. Bioinformatics 38, 1816–1822 (2022).

37. Lei Chen, Z.Z., Junhao Su, Xian Yu, Angel On Ki Wong, Jingcheng Zhang, Yan-Lam Lee, Ruibang Luo A deep-learning method for long-read tumor-only somatic small variant calling. bioRxiv (2025).

38. Benjamin, D. et al. Calling somatic SNVs and indels with Mutect2. BioRxiv, 861054 (2019).

