## Supplementary Information for "Clair-Mosaic: A deep-learning method for long-read mosaic small variant calling"

### Supplementary Figures

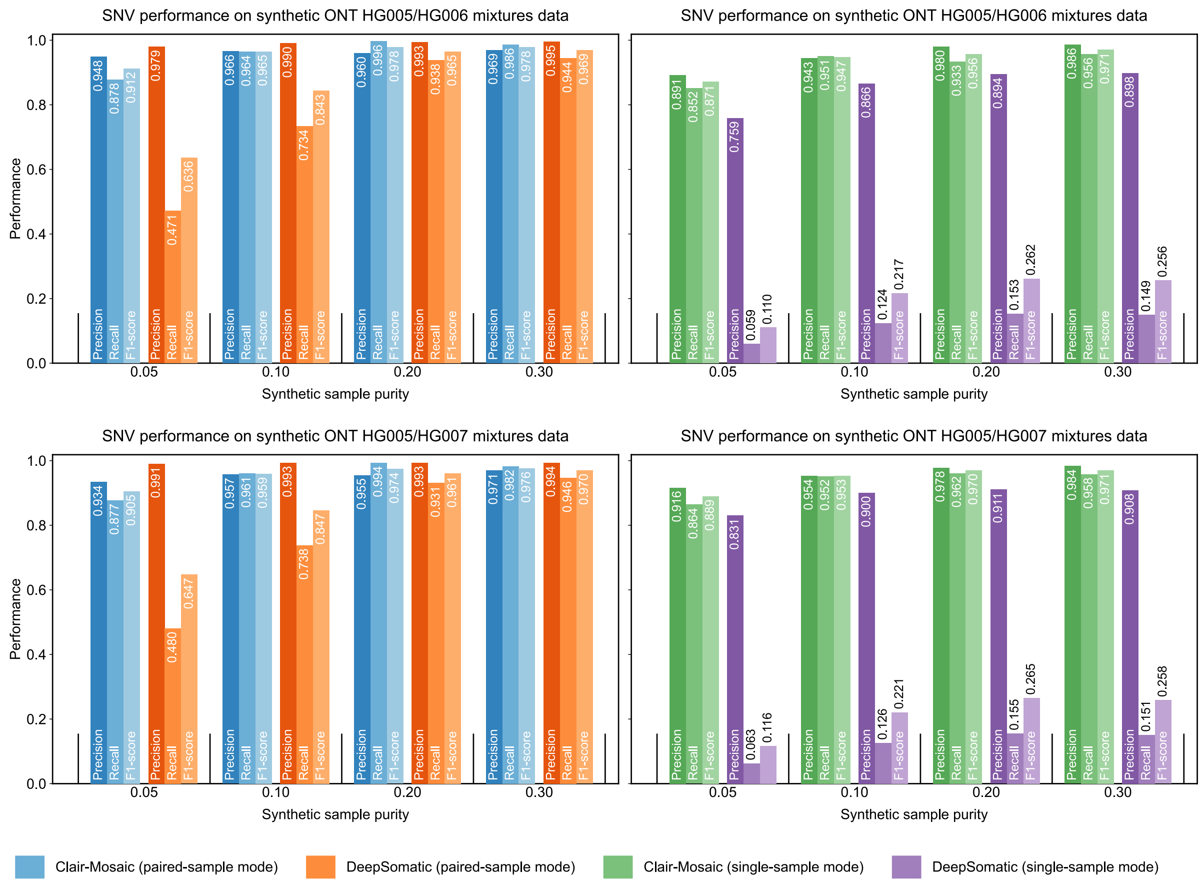

**Supplementary Figure 1. SNV performance at different sample purities in HG005/HG006 and HG005/HG007 mixtures.**

The figure illustrates the SNV performance at different sample purities in HG005/HG006 and HG005/HG007 input/control synthetic dataset mixtures.

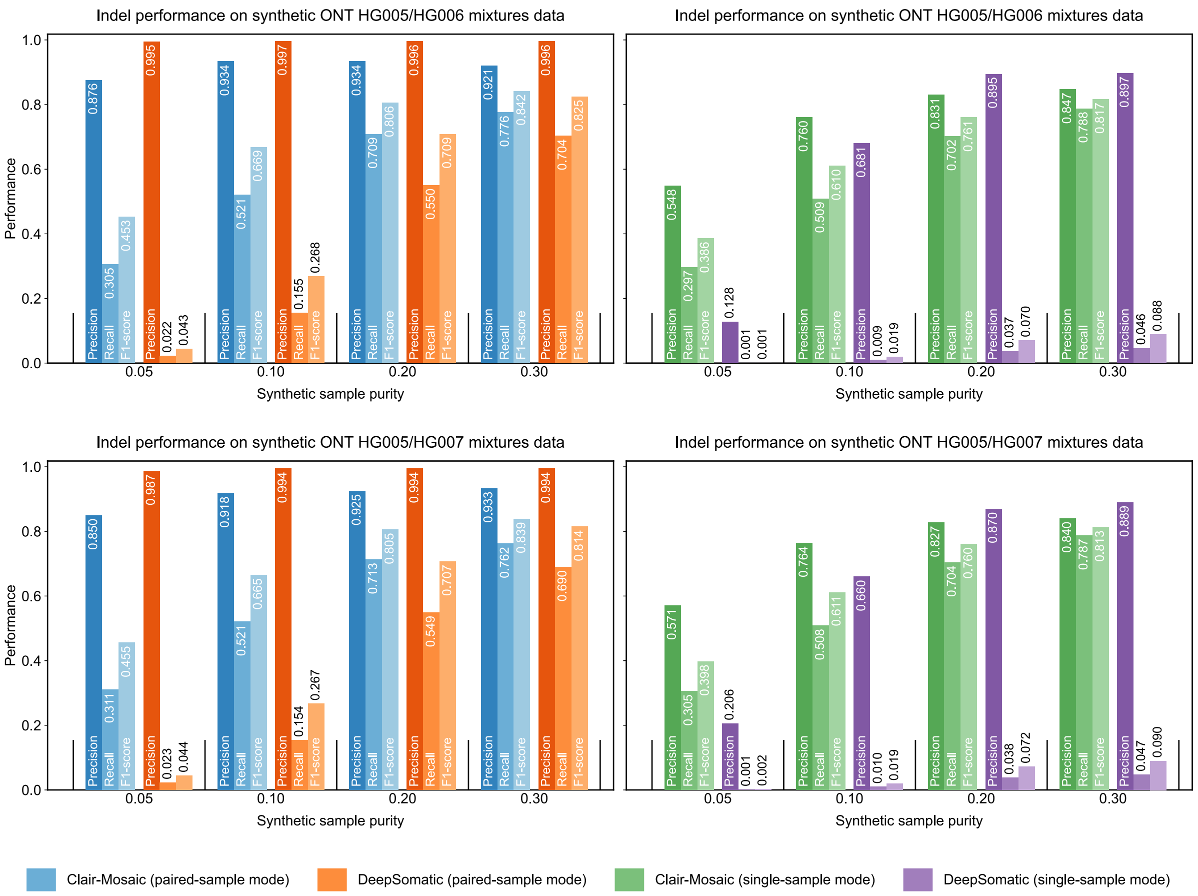

**Supplementary Figure 2. Indel performance at different sample purities in HG005/HG006 and HG005/HG007 mixtures.**

The figure illustrates the Indel performance at different sample purities in HG005/HG006 and HG005/HG007 input/control synthetic dataset mixtures.

### Supplementary Tables

**Supplementary Table 1. Summary of datasets used for model training and benchmarking.**

A summary of sequencing data in ONT, PacBio, and Illumina used for model training and benchmarking.

| Platform | Sample | Reference | Aligner | Coverage | Source | Chemistry /Instrument | Used for data synthesis for model training | Used for benchmarking on synthetic datasets | Used for benchmarking on real samples |
|---|---|---|---|---|---|---|---|---|---|
| ONT | HG001 | GRCh38 | Minimap2 | 78.89 | ONT EPI2ME Labs | R10.4.1 | ✓ | | ✓ |
| | HG002 | GRCh38 | Minimap2 | 91.18 | ONT EPI2ME Labs | R10.4.1 | ✓ | ✓ | ✓ |
| | HG003 | GRCh38 | Minimap2 | 72.51 | ONT EPI2ME Labs | R10.4.1 | | ✓ | ✓ |
| | HG004 | GRCh38 | Minimap2 | 58.84 | ONT EPI2ME Labs | R10.4.1 | | ✓ | ✓ |
| | HG005 | GRCh38 | Minimap2 | 55.00 | ONT EPI2ME Labs | R10.4.1 | | ✓ | ✓ |
| | HG006 | GRCh38 | Minimap2 | 54.89 | ONT EPI2ME Labs | R10.4.1 | | ✓ | ✓ |
| | HG007 | GRCh38 | Minimap2 | 65.95 | ONT EPI2ME Labs | R10.4.1 | | ✓ | ✓ |
| PacBio | HG002 | GRCh38 | pbmm2 | 90.28 | PacBio | HiFi Revio | | ✓ | ✓ |
| | HG003 | GRCh38 | pbmm2 | 61.15 | PacBio | HiFi Revio | ✓ | ✓ | |
| | HG004 | GRCh38 | pbmm2 | 62.67 | PacBio | HiFi Revio | ✓ | ✓ | |
| Illumina | HG002 | GRCh38 | BWA-MEM | 49.82 | Google Health Center | NovaSeq 6000 | | ✓ | ✓ |
| | HG003 | GRCh38 | BWA-MEM | 47.38 | Google Health Center | NovaSeq 6000 | ✓ | ✓ | |
| | HG004 | GRCh38 | BWA-MEM | 46.36 | Google Health Center | NovaSeq 6000 | ✓ | ✓ | |
| | HG003 | GRCh38 | BWA-MEM | 43.77 | Google Health Center | HiseqX | ✓ | | |
| | HG004 | GRCh38 | BWA-MEM | 42.13 | Google Health Center | HiseqX | ✓ | | |

**Supplementary Table 2. ONT SNV result with multiple sample purities in synthetic datasets.**

Benchmark results of multiple sample purities of different callers using **(a)** HG002/HG003, **(b)** HG002/HG004, **(c)** HG005/HG006, and **(d)** HG005/HG007 mixtures.

**a**

| Dataset | Mode | Caller | Coverage | Sample purity | Variant type | Precision | Recall | F1-score | TP | FP | FN |
|---|---|---|---|---|---|---|---|---|---|---|---|
| ONT HG002/HG003 mixtures | Paired-sample mode | Clair-Mosaic | 50x/25x | 0.05 | SNV | 0.8404 | 0.8766 | 0.8581 | 260,647 | 49,511 | 36,692 |
| | | | | | Indel | 0.6939 | 0.2159 | 0.3293 | 10,917 | 4,815 | 39,648 |
| | | | | 0.10 | SNV | 0.8924 | 0.9734 | 0.9311 | 527,901 | 63,647 | 14,421 |
| | | | | | Indel | 0.8383 | 0.4086 | 0.5495 | 29,353 | 5,660 | 42,477 |
| | | | | 0.20 | SNV | 0.9418 | 0.9724 | 0.9569 | 631,512 | 39,016 | 17,933 |
| | | | | | Indel | 0.8650 | 0.5986 | 0.7076 | 49,907 | 7,789 | 33,462 |
| | | | | 0.30 | SNV | 0.9613 | 0.9662 | 0.9638 | 634,136 | 25,515 | 22,168 |
| | | | | | Indel | 0.8954 | 0.6513 | 0.7541 | 55,468 | 6,481 | 29,694 |
| | | DeepSomatic | 50x/25x | 0.05 | SNV | 0.9885 | 0.4028 | 0.5723 | 119,758 | 1,392 | 177,581 |
| | | | | | Indel | 0.9855 | 0.0188 | 0.0369 | 951 | 14 | 49,614 |
| | | | | 0.10 | SNV | 0.9948 | 0.6436 | 0.7816 | 349,050 | 1,825 | 193,272 |
| | | | | | Indel | 0.9965 | 0.1099 | 0.1979 | 7,892 | 28 | 63,938 |
| | | | | 0.20 | SNV | 0.9956 | 0.8919 | 0.9409 | 579,251 | 2,562 | 70,194 |
| | | | | | Indel | 0.9954 | 0.3959 | 0.5665 | 33,008 | 151 | 50,361 |
| | | | | 0.30 | SNV | 0.9965 | 0.9221 | 0.9578 | 605,181 | 2,144 | 51,123 |
| | | | | | Indel | 0.9940 | 0.5822 | 0.7343 | 49,580 | 300 | 35,582 |
| | Single-sample mode | Clair-Mosaic | 50x | 0.05 | SNV | 0.8862 | 0.7748 | 0.8268 | 230,385 | 29,596 | 66,954 |
| | | | | | Indel | 0.4403 | 0.2274 | 0.2999 | 11,500 | 14,616 | 39,065 |
| | | | | 0.10 | SNV | 0.9416 | 0.8937 | 0.9170 | 484,667 | 30,051 | 57,655 |
| | | | | | Indel | 0.6593 | 0.3995 | 0.4975 | 28,698 | 14,832 | 43,132 |
| | | | | 0.20 | SNV | 0.9790 | 0.8915 | 0.9332 | 578,966 | 12,394 | 70,479 |
| | | | | | Indel | 0.7599 | 0.5762 | 0.6554 | 48,038 | 15,181 | 35,331 |
| | | | | 0.30 | SNV | 0.9856 | 0.9136 | 0.9482 | 599,610 | 8,778 | 56,694 |
| | | | | | Indel | 0.7863 | 0.6536 | 0.7139 | 55,664 | 15,127 | 29,498 |
| | | DeepSomatic | 50x | 0.05 | SNV | 0.7704 | 0.1374 | 0.2331 | 40,841 | 12,175 | 256,498 |
| | | | | | Indel | 0.1345 | 0.0020 | 0.0039 | 101 | 650 | 50,464 |
| | | | | 0.10 | SNV | 0.8782 | 0.1573 | 0.2668 | 85,318 | 11,835 | 457,004 |
| | | | | | Indel | 0.5164 | 0.0097 | 0.0190 | 694 | 650 | 71,136 |
| | | | | 0.20 | SNV | 0.9086 | 0.1749 | 0.2934 | 113,600 | 11,425 | 535,845 |
| | | | | | Indel | 0.8002 | 0.0316 | 0.0608 | 2,635 | 658 | 80,734 |
| | | | | 0.30 | SNV | 0.9207 | 0.1758 | 0.2952 | 115,351 | 9,936 | 540,953 |
| | | | | | Indel | 0.8405 | 0.0440 | 0.0836 | 3,746 | 711 | 81,416 |

**b**

| Dataset | Mode | Caller | Coverage | Sample purity | Variant type | Precision | Recall | F1-score | TP | FP | FN |
|---|---|---|---|---|---|---|---|---|---|---|---|
| ONT HG002/HG004 mixtures | Paired-sample mode | Clair-Mosaic | 50x/25x | 0.05 | SNV | 0.8284 | 0.9118 | 0.8681 | 228,308 | 47,306 | 22,086 |
| | | | | | Indel | 0.7290 | 0.2127 | 0.3293 | 9,848 | 3,660 | 36,461 |
| | | | | 0.10 | SNV | 0.8868 | 0.9828 | 0.9324 | 507,407 | 64,757 | 8,857 |
| | | | | | Indel | 0.8587 | 0.4155 | 0.5600 | 28,632 | 4,711 | 40,284 |
| | | | | 0.20 | SNV | 0.9490 | 0.9833 | 0.9659 | 627,265 | 33,701 | 10,649 |
| | | | | | Indel | 0.8750 | 0.6016 | 0.7130 | 49,307 | 7,047 | 32,654 |
| | | | | 0.30 | SNV | 0.9643 | 0.9721 | 0.9682 | 627,834 | 23,222 | 18,003 |
| | | | | | Indel | 0.8992 | 0.6555 | 0.7582 | 54,919 | 6,159 | 28,866 |
| | | DeepSomatic | 50x/25x | 0.05 | SNV | 0.9951 | 0.3994 | 0.5700 | 100,000 | 492 | 150,394 |
| | | | | | Indel | 0.9916 | 0.0152 | 0.0300 | 706 | 6 | 45,603 |
| | | | | 0.10 | SNV | 0.9959 | 0.6466 | 0.7841 | 333,815 | 1,389 | 182,449 |
| | | | | | Indel | 0.9961 | 0.1034 | 0.1874 | 7,128 | 28 | 61,788 |
| | | | | 0.20 | SNV | 0.9958 | 0.9186 | 0.9556 | 585,959 | 2,495 | 51,955 |
| | | | | | Indel | 0.9963 | 0.3981 | 0.5689 | 32,626 | 121 | 49,335 |
| | | | | 0.30 | SNV | 0.9960 | 0.9298 | 0.9618 | 600,523 | 2,398 | 45,314 |
| | | | | | Indel | 0.9944 | 0.5919 | 0.7421 | 49,593 | 277 | 34,192 |
| | Single-sample mode | Clair-Mosaic | 50x | 0.05 | SNV | 0.8711 | 0.7531 | 0.8078 | 188,564 | 27,900 | 61,830 |
| | | | | | Indel | 0.3688 | 0.1966 | 0.2565 | 9,104 | 15,583 | 37,205 |
| | | | | 0.10 | SNV | 0.9397 | 0.8861 | 0.9121 | 457,455 | 29,362 | 58,809 |
| | | | | | Indel | 0.6292 | 0.3867 | 0.4790 | 26,649 | 15,703 | 42,267 |
| | | | | 0.20 | SNV | 0.9786 | 0.9215 | 0.9492 | 587,818 | 12,840 | 50,096 |
| | | | | | Indel | 0.7472 | 0.5696 | 0.6464 | 46,681 | 15,796 | 35,280 |
| | | | | 0.30 | SNV | 0.9830 | 0.9451 | 0.9637 | 610,390 | 10,583 | 35,447 |
| | | | | | Indel | 0.7775 | 0.6518 | 0.7091 | 54,614 | 15,628 | 29,171 |
| | | DeepSomatic | 50x | 0.05 | SNV | 0.7693 | 0.1327 | 0.2263 | 33,222 | 9,963 | 217,172 |
| | | | | | Indel | 0.0996 | 0.0016 | 0.0031 | 73 | 660 | 46,236 |
| | | | | 0.10 | SNV | 0.8852 | 0.1521 | 0.2596 | 78,528 | 10,181 | 437,736 |
| | | | | | Indel | 0.4902 | 0.0094 | 0.0184 | 647 | 673 | 68,269 |
| | | | | 0.20 | SNV | 0.9158 | 0.1695 | 0.2861 | 108,136 | 9,945 | 529,778 |
| | | | | | Indel | 0.7941 | 0.0310 | 0.0596 | 2,538 | 658 | 79,423 |
| | | | | 0.30 | SNV | 0.9181 | 0.1702 | 0.2872 | 109,929 | 9,810 | 535,908 |
| | | | | | Indel | 0.8346 | 0.0430 | 0.0818 | 3,603 | 714 | 80,182 |

**c**

| Dataset | Mode | Caller | Coverage | Sample purity | Variant type | Precision | Recall | F1-score | TP | FP | FN |
|---|---|---|---|---|---|---|---|---|---|---|---|
| ONT HG005/HG006 mixtures | Paired-sample mode | Clair-Mosaic | 50x/25x | 0.05 | SNV | 0.9481 | 0.8778 | 0.9116 | 238,858 | 13,084 | 33,265 |
| | | | | | Indel | 0.8762 | 0.3052 | 0.4527 | 11,403 | 1,611 | 25,965 |
| | | | | 0.10 | SNV | 0.9658 | 0.9638 | 0.9648 | 492,386 | 17,438 | 18,483 |
| | | | | | Indel | 0.9336 | 0.5207 | 0.6686 | 30,105 | 2,142 | 27,706 |
| | | | | 0.20 | SNV | 0.9595 | 0.9961 | 0.9775 | 579,353 | 24,482 | 2,244 |
| | | | | | Indel | 0.9337 | 0.7087 | 0.8058 | 46,322 | 3,291 | 19,041 |
| | | | | 0.30 | SNV | 0.9692 | 0.9859 | 0.9775 | 575,474 | 18,302 | 8,205 |
| | | | | | Indel | 0.9209 | 0.7758 | 0.8421 | 51,124 | 4,394 | 14,778 |
| | | DeepSomatic | 50x/25x | 0.05 | SNV | 0.9793 | 0.4714 | 0.6364 | 128,278 | 2,714 | 143,845 |
| | | | | | Indel | 0.9952 | 0.0222 | 0.0435 | 831 | 4 | 36,537 |
| | | | | 0.10 | SNV | 0.9903 | 0.7335 | 0.8428 | 374,714 | 3,665 | 136,155 |
| | | | | | Indel | 0.9967 | 0.1549 | 0.2681 | 8,955 | 30 | 48,856 |
| | | | | 0.20 | SNV | 0.9934 | 0.9375 | 0.9646 | 545,236 | 3,650 | 36,361 |
| | | | | | Indel | 0.9957 | 0.5505 | 0.7090 | 35,981 | 156 | 29,382 |
| | | | | 0.30 | SNV | 0.9948 | 0.9438 | 0.9686 | 550,884 | 2,891 | 32,795 |
| | | | | | Indel | 0.9957 | 0.7035 | 0.8245 | 46,360 | 198 | 19,542 |
| | Single-sample mode | Clair-Mosaic | 50x | 0.05 | SNV | 0.8909 | 0.8516 | 0.8708 | 231,738 | 28,369 | 40,385 |
| | | | | | Indel | 0.5484 | 0.2972 | 0.3855 | 11,106 | 9,144 | 26,262 |
| | | | | 0.10 | SNV | 0.9434 | 0.9511 | 0.9472 | 485,872 | 29,174 | 24,997 |
| | | | | | Indel | 0.7605 | 0.5093 | 0.6101 | 29,445 | 9,275 | 28,366 |
| | | | | 0.20 | SNV | 0.9796 | 0.9332 | 0.9558 | 542,738 | 11,293 | 38,859 |
| | | | | | Indel | 0.8307 | 0.7017 | 0.7607 | 45,864 | 9,349 | 19,499 |
| | | | | 0.30 | SNV | 0.9861 | 0.9563 | 0.9710 | 558,158 | 7,869 | 25,521 |
| | | | | | Indel | 0.8472 | 0.7880 | 0.8165 | 51,929 | 9,366 | 13,973 |
| | | DeepSomatic | 50x | 0.05 | SNV | 0.7587 | 0.0593 | 0.1100 | 34,621 | 11,008 | 549,289 |
| | | | | | Indel | 0.1283 | 0.0007 | 0.0013 | 44 | 299 | 66,243 |
| | | | | 0.10 | SNV | 0.8656 | 0.1239 | 0.2167 | 72,320 | 11,233 | 511,590 |
| | | | | | Indel | 0.6807 | 0.0095 | 0.0187 | 629 | 295 | 65,658 |
| | | | | 0.20 | SNV | 0.8945 | 0.1532 | 0.2616 | 89,092 | 10,510 | 492,505 |
| | | | | | Indel | 0.8948 | 0.0366 | 0.0703 | 2,391 | 281 | 62,972 |
| | | | | 0.30 | SNV | 0.8982 | 0.1493 | 0.2561 | 87,157 | 9,877 | 496,522 |
| | | | | | Indel | 0.8973 | 0.0464 | 0.0883 | 3,059 | 350 | 62,843 |

**d**

| Dataset | Mode | Caller | Coverage | Sample purity | Variant type | Precision | Recall | F1-score | TP | FP | FN |
|---|---|---|---|---|---|---|---|---|---|---|---|
| ONT HG005/HG007 mixtures | Paired-sample mode | Clair-Mosaic | 50x/25x | 0.05 | SNV | 0.9343 | 0.8771 | 0.9048 | 239,872 | 16,877 | 33,604 |
| | | | | | Indel | 0.8497 | 0.3109 | 0.4552 | 11,648 | 2,061 | 25,823 |
| | | | | 0.10 | SNV | 0.9571 | 0.9610 | 0.9590 | 484,682 | 21,742 | 19,663 |
| | | | | | Indel | 0.9181 | 0.5210 | 0.6647 | 29,743 | 2,652 | 27,349 |
| | | | | 0.20 | SNV | 0.9551 | 0.9936 | 0.9740 | 566,555 | 26,620 | 3,655 |
| | | | | | Indel | 0.9251 | 0.7127 | 0.8051 | 45,858 | 3,714 | 18,490 |
| | | | | 0.30 | SNV | 0.9706 | 0.9820 | 0.9763 | 561,626 | 17,035 | 10,275 |
| | | | | | Indel | 0.9333 | 0.7617 | 0.8388 | 49,391 | 3,532 | 15,454 |
| | | DeepSomatic | 50x/25x | 0.05 | SNV | 0.9908 | 0.4805 | 0.6472 | 131,414 | 1,219 | 142,062 |
| | | | | | Indel | 0.9872 | 0.0227 | 0.0443 | 849 | 11 | 36,622 |
| | | | | 0.10 | SNV | 0.9934 | 0.7380 | 0.8468 | 372,185 | 2,480 | 132,160 |
| | | | | | Indel | 0.9943 | 0.1540 | 0.2667 | 8,793 | 50 | 48,299 |
| | | | | 0.20 | SNV | 0.9934 | 0.9311 | 0.9613 | 530,951 | 3,512 | 39,259 |
| | | | | | Indel | 0.9942 | 0.5487 | 0.7071 | 35,307 | 206 | 29,041 |
| | | | | 0.30 | SNV | 0.9940 | 0.9464 | 0.9696 | 541,226 | 3,272 | 30,675 |
| | | | | | Indel | 0.9940 | 0.6898 | 0.8144 | 44,728 | 268 | 20,117 |
| | Single-sample mode | Clair-Mosaic | 50x | 0.05 | SNV | 0.9163 | 0.8641 | 0.8894 | 236,307 | 21,592 | 37,169 |
| | | | | | Indel | 0.5710 | 0.3053 | 0.3978 | 11,439 | 8,596 | 26,032 |
| | | | | 0.10 | SNV | 0.9535 | 0.9519 | 0.9527 | 480,107 | 23,403 | 24,238 |
| | | | | | Indel | 0.7645 | 0.5083 | 0.6106 | 29,019 | 8,937 | 28,073 |
| | | | | 0.20 | SNV | 0.9782 | 0.9615 | 0.9698 | 548,272 | 12,219 | 21,938 |
| | | | | | Indel | 0.8268 | 0.7041 | 0.7605 | 45,309 | 9,491 | 19,039 |
| | | | | 0.30 | SNV | 0.9835 | 0.9578 | 0.9705 | 547,761 | 9,209 | 24,140 |
| | | | | | Indel | 0.8403 | 0.7872 | 0.8129 | 51,044 | 9,699 | 13,801 |
| | | DeepSomatic | 50x | 0.05 | SNV | 0.8307 | 0.0626 | 0.1164 | 35,812 | 7,297 | 536,288 |
| | | | | | Indel | 0.2056 | 0.0011 | 0.0022 | 73 | 282 | 65,170 |
| | | | | 0.10 | SNV | 0.9005 | 0.1258 | 0.2208 | 71,972 | 7,956 | 500,128 |
| | | | | | Indel | 0.6597 | 0.0096 | 0.0190 | 628 | 324 | 64,615 |
| | | | | 0.20 | SNV | 0.9106 | 0.1552 | 0.2652 | 88,478 | 8,684 | 481,732 |
| | | | | | Indel | 0.8698 | 0.0377 | 0.0722 | 2,424 | 363 | 61,924 |
| | | | | 0.30 | SNV | 0.9084 | 0.1506 | 0.2584 | 86,145 | 8,690 | 485,756 |
| | | | | | Indel | 0.8895 | 0.0473 | 0.0898 | 3,067 | 381 | 61,778 |

**Supplementary Table 3. ONT SNV result with multiple allelic fractions in synthetic datasets.**

Benchmark results of Clair-Mosaic with multiple allelic fractions in HG002/HG003 synthetic dataset mixtures.

| Dataset | Mode | Caller | Coverage | AF range | Variant type | Precision | Recall | F1-score | TP | FP | FN |
|---|---|---|---|---|---|---|---|---|---|---|---|
| ONT HG002/HG003 mixtures | Paired-sample mode | Clair-Mosaic | 50x/25x | 0.05-0.1 | SNV | 0.7285 | 0.8818 | 0.7978 | 383,410 | 142,924 | 51,391 |
| | | | | | Indel | 0.6520 | 0.3360 | 0.4435 | 16,745 | 8,939 | 33,085 |
| | | | | 0.1-0.2 | SNV | 0.9404 | 0.9815 | 0.9605 | 743,569 | 47,153 | 13,986 |
| | | | | | Indel | 0.8157 | 0.4904 | 0.6126 | 54,540 | 12,320 | 56,668 |
| | | | | 0.2-0.3 | SNV | 0.9900 | 0.9944 | 0.9922 | 589,230 | 5,937 | 3,316 |
| | | | | | Indel | 0.9103 | 0.5708 | 0.7016 | 50,028 | 4,928 | 37,622 |
| | | DeepSomatic | 50x/25x | 0.05-0.1 | SNV | 0.9879 | 0.3259 | 0.4902 | 141,718 | 1,739 | 293,083 |
| | | | | | Indel | 1.0000 | 0.0018 | 0.0036 | 89 | 0 | 49,741 |
| | | | | 0.1-0.2 | SNV | 0.9947 | 0.8294 | 0.9045 | 628,322 | 3,374 | 129,233 |
| | | | | | Indel | 0.9946 | 0.2254 | 0.3676 | 25,071 | 135 | 86,137 |
| | | | | 0.2-0.3 | SNV | 0.9957 | 0.9708 | 0.9831 | 575,245 | 2,484 | 17,301 |
| | | | | | Indel | 0.9947 | 0.4804 | 0.6479 | 42,104 | 225 | 45,546 |
| | Single-sample mode | Clair-Mosaic | 50x | 0.05-0.1 | SNV | 0.8914 | 0.7511 | 0.8152 | 326,573 | 39,794 | 108,228 |
| | | | | | Indel | 0.4571 | 0.3826 | 0.4165 | 19,064 | 22,639 | 30,766 |
| | | | | 0.1-0.2 | SNV | 0.9460 | 0.9685 | 0.9572 | 733,720 | 41,859 | 23,835 |
| | | | | | Indel | 0.4797 | 0.5472 | 0.5112 | 60,848 | 66,007 | 50,360 |
| | | | | 0.2-0.3 | SNV | 0.9695 | 0.9984 | 0.9837 | 591,578 | 18,616 | 968 |
| | | | | | Indel | 0.3808 | 0.6822 | 0.4888 | 59,792 | 97,217 | 27,858 |
| | | DeepSomatic | 50x | 0.05-0.1 | SNV | 0.6860 | 0.1359 | 0.2269 | 59,093 | 27,052 | 375,708 |
| | | | | | Indel | 1.0000 | 0.0002 | 0.0004 | 11 | 0 | 49,819 |
| | | | | 0.1-0.2 | SNV | 0.9240 | 0.1688 | 0.2854 | 127,867 | 10,522 | 629,688 |
| | | | | | Indel | 0.7017 | 0.0174 | 0.0339 | 1,931 | 821 | 109,277 |
| | | | | 0.2-0.3 | SNV | 0.9685 | 0.1786 | 0.3016 | 105,815 | 3,443 | 486,731 |
| | | | | | Indel | 0.8077 | 0.0369 | 0.0706 | 3,235 | 770 | 84,415 |

**Supplementary Table 4. Benchmark results across different genomic contexts.**

Benchmark results of Clair-Mosaic in HG002/HG003 synthetic dataset mixtures across different genomic contexts, defined as GIAB stratifications, in **(a)** paired-sample mode and **(b)** single-sample mode.

**a**

| Dataset | Mode | Caller | Coverage | Stratification type | Stratification subtype | Variant type | Precision | Recall | F1-score | TP | FP | FN |
|---|---|---|---|---|---|---|---|---|---|---|---|---|
| ONT HG002/HG003 mixtures | Paired-sample Mode | Clair-Mosaic | 50x/25x | Low complexity | Homopol_4-6bp | SNV | 0.9106 | 0.9003 | 0.9054 | 481,280 | 47,258 | 53,305 |
| | | | | | | Indel | 0.8674 | 0.5717 | 0.6892 | 47,932 | 7,326 | 35,903 |
| | | | | | Homopol_7-11bp | SNV | 0.6265 | 0.9006 | 0.7389 | 41,248 | 24,594 | 4,552 |
| | | | | | | Indel | 0.6735 | 0.2495 | 0.3641 | 17,803 | 8,632 | 53,550 |
| | | | | | Homopol_ge12bp | SNV | 0.2951 | 0.8497 | 0.4380 | 5,115 | 12,219 | 905 |
| | | | | | | Indel | 0.2313 | 0.0122 | 0.0231 | 639 | 2,124 | 51,904 |
| | | | | | Imp_Homopol_ge11bp | SNV | 0.4588 | 0.8929 | 0.6062 | 20,530 | 24,215 | 2,462 |
| | | | | | | Indel | 0.6085 | 0.0895 | 0.1560 | 7,232 | 4,653 | 73,596 |
| | | | | | TR_le50bp | SNV | 0.8534 | 0.8904 | 0.8715 | 19,391 | 3,330 | 2,387 |
| | | | | | | Indel | 0.8261 | 0.6000 | 0.6951 | 17,800 | 3,747 | 11,866 |
| | | | | | TR51-220bp | SNV | 0.8096 | 0.8524 | 0.8305 | 15,007 | 3,529 | 2,598 |
| | | | | | | Indel | 0.8439 | 0.6442 | 0.7307 | 9,651 | 1,785 | 5,330 |
| | | | | | TR201-10kbp | SNV | 0.9067 | 0.9065 | 0.9066 | 6,822 | 702 | 704 |
| | | | | | | Indel | 0.9422 | 0.7746 | 0.8502 | 1,808 | 111 | 526 |
| | | | | | TR_ge101bp | SNV | 0.8762 | 0.8876 | 0.8819 | 12,812 | 1,810 | 1,622 |
| | | | | | | Indel | 0.8805 | 0.7041 | 0.7825 | 5,327 | 723 | 2,239 |
| | | | | | TR_and_Homopol | SNV | 0.7040 | 0.8835 | 0.7836 | 95,050 | 39,970 | 12,536 |
| | | | | | | Indel | 0.7448 | 0.2983 | 0.4260 | 51,897 | 17,785 | 122,083 |
| | | | | Segmental duplications | SegDups | SNV | 0.8371 | 0.8662 | 0.8514 | 56,747 | 11,042 | 8,766 |
| | | | | | | Indel | 0.8207 | 0.7157 | 0.7647 | 3,223 | 704 | 1,280 |
| | | | | | SegDups_gt10kb | SNV | 0.8234 | 0.8619 | 0.8422 | 47,648 | 10,218 | 7,637 |
| | | | | | | Indel | 0.8061 | 0.7089 | 0.7544 | 2,628 | 632 | 1,079 |
| | | | | Low mappability | LowMap | SNV | 0.8961 | 0.9043 | 0.9002 | 107,918 | 12,516 | 11,419 |
| | | | | | | Indel | 0.8546 | 0.6926 | 0.7651 | 3,761 | 640 | 1,669 |
| | | | | Other difficult regions | L1H | SNV | 0.9718 | 0.9327 | 0.9519 | 3,452 | 100 | 249 |
| | | | | | | Indel | 0.9474 | 0.6716 | 0.7860 | 90 | 5 | 44 |
| | | | | Functional regions | CDS | SNV | 0.9211 | 0.9001 | 0.9104 | 11,959 | 1,025 | 1,328 |
| | | | | | | Indel | 0.8673 | 0.6753 | 0.7593 | 183 | 28 | 88 |
| | | DeepSomatic | 50x/25x | Low complexity | Homopol_4-6bp | SNV | 0.9948 | 0.7807 | 0.8748 | 417,326 | 2,198 | 117,259 |
| | | | | | | Indel | 0.9945 | 0.3723 | 0.5418 | 31,211 | 174 | 52,624 |
| | | | | | Homopol_7-11bp | SNV | 0.9964 | 0.5617 | 0.7184 | 25,728 | 93 | 20,072 |
| | | | | | | Indel | 0.9905 | 0.1846 | 0.3112 | 13,172 | 127 | 58,181 |
| | | | | | Homopol_ge12bp | SNV | 0.9908 | 0.2681 | 0.4220 | 1,614 | 15 | 4,406 |
| | | | | | | Indel | 0.9617 | 0.0043 | 0.0086 | 226 | 9 | 52,300 |
| | | | | | Imp_Homopol_ge11bp | SNV | 0.9978 | 0.4346 | 0.6055 | 9,992 | 22 | 13,000 |
| | | | | | | Indel | 0.9890 | 0.0578 | 0.1092 | 4,673 | 52 | 76,155 |
| | | | | | TR_le50bp | SNV | 0.9938 | 0.7250 | 0.8384 | 15,789 | 98 | 5,989 |
| | | | | | | Indel | 0.9972 | 0.3334 | 0.4997 | 9,891 | 28 | 19,775 |
| | | | | | TR51-220bp | SNV | 0.9936 | 0.7077 | 0.8266 | 12,459 | 80 | 5,146 |
| | | | | | | Indel | 0.9950 | 0.3347 | 0.5009 | 5,014 | 25 | 9,967 |
| | | | | | TR201-10kbp | SNV | 0.9921 | 0.7378 | 0.8463 | 5,553 | 44 | 1,973 |
| | | | | | | Indel | 0.9951 | 0.4340 | 0.6044 | 1,013 | 5 | 1,321 |
| | | | | | TR_ge101bp | SNV | 0.9926 | 0.7188 | 0.8338 | 10,375 | 77 | 4,059 |
| | | | | | | Indel | 0.9954 | 0.3719 | 0.5415 | 2,814 | 13 | 4,752 |
| | | | | | TR_and_Homopol | SNV | 0.9965 | 0.5989 | 0.7481 | 64,431 | 227 | 43,155 |
| | | | | | | Indel | 0.9935 | 0.1813 | 0.3066 | 31,539 | 207 | 142,441 |
| | | | | Segmental duplications | SegDups | SNV | 0.9881 | 0.6748 | 0.8019 | 44,205 | 534 | 21,308 |
| | | | | | | Indel | 0.9791 | 0.4275 | 0.5951 | 1,925 | 41 | 2,578 |
| | | | | | SegDups_gt10kb | SNV | 0.9897 | 0.6586 | 0.7909 | 36,413 | 378 | 18,872 |
| | | | | | | Indel | 0.9782 | 0.4233 | 0.5908 | 1,569 | 35 | 2,138 |
| | | | | Low mappability | LowMap | SNV | 0.9947 | 0.6681 | 0.7993 | 79,728 | 421 | 39,609 |
| | | | | | | Indel | 0.9858 | 0.4092 | 0.5783 | 2,222 | 32 | 3,208 |
| | | | | Other difficult regions | L1H | SNV | 1.0000 | 0.5782 | 0.7328 | 2,140 | 0 | 1,561 |
| | | | | | | Indel | 1.0000 | 0.3488 | 0.5172 | 45 | 0 | 84 |
| | | | | Functional regions | CDS | SNV | 0.9869 | 0.7983 | 0.8826 | 10,607 | 141 | 2,680 |
| | | | | | | Indel | 1.0000 | 0.3727 | 0.5430 | 101 | 0 | 170 |

**b**

| Dataset | Mode | Caller | Coverage | Stratification type | Stratification subtype | Variant type | Precision | Recall | F1-score | TP | FP | FN |
|---|---|---|---|---|---|---|---|---|---|---|---|---|
| ONT HG002/HG003 mixtures | Single-sample Mode | Clair-Mosaic | 50x | Low complexity | Homopol_4-6bp | SNV | 0.9571 | 0.8128 | 0.8791 | 434,511 | 19,496 | 100,074 |
| | | | | | | Indel | 0.7295 | 0.5440 | 0.6232 | 45,603 | 16,907 | 38,232 |
| | | | | | Homopol_7-11bp | SNV | 0.9183 | 0.6310 | 0.7480 | 28,898 | 2,572 | 16,902 |
| | | | | | | Indel | 0.3177 | 0.3053 | 0.3114 | 21,785 | 46,792 | 49,568 |
| | | | | | Homopol_ge12bp | SNV | 0.6714 | 0.3478 | 0.4583 | 2,094 | 1,025 | 3,926 |
| | | | | | | Indel | 0.0732 | 0.1405 | 0.0962 | 7,380 | 93,498 | 45,163 |
| | | | | | Imp_Homopol_ge11bp | SNV | 0.8720 | 0.5677 | 0.6877 | 13,053 | 1,916 | 9,939 |
| | | | | | | Indel | 0.1483 | 0.1547 | 0.1514 | 12,501 | 71,796 | 68,327 |
| | | | | | TR_le50bp | SNV | 0.9246 | 0.7307 | 0.8163 | 15,913 | 1,297 | 5,865 |
| | | | | | | Indel | 0.6108 | 0.6820 | 0.6444 | 20,232 | 12,892 | 9,434 |
| | | | | | TR51-220bp | SNV | 0.9073 | 0.7289 | 0.8084 | 12,833 | 1,311 | 4,772 |
| | | | | | | Indel | 0.6416 | 0.7030 | 0.6709 | 10,531 | 5,882 | 4,450 |
| | | | | | TR201-10kbp | SNV | 0.9456 | 0.8286 | 0.8832 | 6,236 | 359 | 1,290 |
| | | | | | | Indel | 0.7934 | 0.8063 | 0.7998 | 1,882 | 490 | 452 |
| | | | | | TR_ge101bp | SNV | 0.9326 | 0.7893 | 0.8550 | 11,393 | 824 | 3,041 |
| | | | | | | Indel | 0.7027 | 0.7367 | 0.7193 | 5,574 | 2,358 | 1,992 |
| | | | | | TR_and_Homopol | SNV | 0.9199 | 0.6753 | 0.7789 | 72,658 | 6,325 | 34,928 |
| | | | | | | Indel | 0.4437 | 0.3403 | 0.3852 | 59,214 | 74,231 | 114,766 |
| | | | | Segmental duplications | SegDups | SNV | 0.8312 | 0.7556 | 0.7916 | 49,502 | 10,050 | 16,011 |
| | | | | | | Indel | 0.5148 | 0.6298 | 0.5665 | 2,836 | 2,673 | 1,667 |
| | | | | | SegDups_gt10kb | SNV | 0.8288 | 0.7475 | 0.7861 | 41,324 | 8,534 | 13,961 |
| | | | | | | Indel | 0.4989 | 0.6383 | 0.5601 | 2,366 | 2,376 | 1,341 |
| | | | | Low mappability | LowMap | SNV | 0.9033 | 0.7856 | 0.8404 | 93,750 | 10,032 | 25,587 |
| | | | | | | Indel | 0.6121 | 0.6692 | 0.6394 | 3,634 | 2,303 | 1,796 |
| | | | | Other difficult regions | L1H | SNV | 0.9951 | 0.7152 | 0.8323 | 2,647 | 13 | 1,054 |
| | | | | | | Indel | 0.8942 | 0.6940 | 0.7815 | 93 | 11 | 41 |
| | | | | Functional regions | CDS | SNV | 0.9080 | 0.8398 | 0.8726 | 11,158 | 1,130 | 2,129 |
| | | | | | | Indel | 0.7441 | 0.5793 | 0.6515 | 157 | 54 | 114 |
| | | DeepSomatic | 50x | Low complexity | Homopol_4-6bp | SNV | 0.8758 | 0.1657 | 0.2787 | 88,600 | 12,559 | 445,985 |
| | | | | | | Indel | 0.6383 | 0.0279 | 0.0535 | 2,340 | 1,326 | 81,495 |
| | | | | | Homopol_7-11bp | SNV | 0.6534 | 0.1461 | 0.2388 | 6,690 | 3,548 | 39,110 |
| | | | | | | Indel | 0.3200 | 0.0055 | 0.0108 | 392 | 833 | 70,961 |
| | | | | | Homopol_ge12bp | SNV | 0.2820 | 0.1249 | 0.1731 | 752 | 1,915 | 5,268 |
| | | | | | | Indel | 0.0576 | 0.0002 | 0.0004 | 11 | 180 | 52,532 |
| | | | | | Imp_Homopol_ge11bp | SNV | 0.5211 | 0.1468 | 0.2291 | 3,376 | 3,102 | 19,616 |
| | | | | | | Indel | 0.4097 | 0.0042 | 0.0083 | 338 | 487 | 80,490 |
| | | | | | TR_le50bp | SNV | 0.7851 | 0.1513 | 0.2537 | 3,295 | 902 | 18,483 |
| | | | | | | Indel | 0.7901 | 0.0249 | 0.0482 | 738 | 196 | 28,928 |
| | | | | | TR51-220bp | SNV | 0.6353 | 0.1588 | 0.2541 | 2,796 | 1,605 | 14,809 |
| | | | | | | Indel | 0.7945 | 0.0253 | 0.0490 | 379 | 98 | 14,602 |
| | | | | | TR201-10kbp | SNV | 0.7825 | 0.1472 | 0.2478 | 1,108 | 308 | 6,418 |
| | | | | | | Indel | 0.8022 | 0.0313 | 0.0602 | 73 | 18 | 2,261 |
| | | | | | TR_ge101bp | SNV | 0.7329 | 0.1472 | 0.2451 | 2,124 | 774 | 12,310 |
| | | | | | | Indel | 0.7946 | 0.0271 | 0.0524 | 205 | 53 | 7,361 |
| | | | | | TR_and_Homopol | SNV | 0.6806 | 0.1504 | 0.2463 | 16,179 | 7,592 | 91,407 |
| | | | | | | Indel | 0.5685 | 0.0105 | 0.0207 | 1,834 | 1,392 | 172,146 |
| | | | | Segmental duplications | SegDups | SNV | 0.6610 | 0.1787 | 0.2814 | 11,708 | 6,004 | 53,805 |
| | | | | | | Indel | 0.5979 | 0.0380 | 0.0714 | 171 | 115 | 4,332 |
| | | | | | SegDups_gt10kb | SNV | 0.6453 | 0.1826 | 0.2846 | 10,094 | 5,549 | 45,191 |
| | | | | | | Indel | 0.6098 | 0.0405 | 0.0759 | 150 | 96 | 3,557 |
| | | | | Low mappability | LowMap | SNV | 0.7634 | 0.1632 | 0.2689 | 19,472 | 6,036 | 99,865 |
| | | | | | | Indel | 0.5319 | 0.0368 | 0.0689 | 200 | 176 | 5,230 |
| | | | | Other difficult regions | L1H | SNV | 0.9306 | 0.1956 | 0.3233 | 724 | 54 | 2,977 |
| | | | | | | Indel | 1.0000 | 0.0310 | 0.0602 | 4 | 0 | 125 |
| | | | | Functional regions | CDS | SNV | 0.8301 | 0.2294 | 0.3595 | 3,048 | 624 | 10,239 |
| | | | | | | Indel | 0.6538 | 0.1255 | 0.2105 | 34 | 18 | 237 |

**Supplementary Table 5. SNV result with multiple sample purities in PacBio synthetic datasets.**

Benchmark results of multiple sample purities of different callers using **(a)** HG002/HG003, **(b)** HG002/HG004 mixtures in the PacBio dataset.

**a**

| Dataset | Mode | Caller | Coverage | Sample purity | Variant type | Precision | Recall | F1-score | TP | FP | FN |
|---|---|---|---|---|---|---|---|---|---|---|---|
| PacBio HG002/HG003 mixtures | Paired-sample mode | Clair-Mosaic | 50x/25x | 0.05 | SNV | 0.9706 | 0.9918 | 0.9811 | 278,136 | 8,431 | 2,304 |
| | | | | | Indel | 0.8350 | 0.1982 | 0.3204 | 9,573 | 1,892 | 38,726 |
| | | | | 0.10 | SNV | 0.9825 | 0.9970 | 0.9897 | 564,658 | 10,049 | 1,689 |
| | | | | | Indel | 0.9285 | 0.4039 | 0.5630 | 30,535 | 2,351 | 45,057 |
| | | | | 0.20 | SNV | 0.9804 | 0.9983 | 0.9893 | 652,345 | 13,040 | 1,083 |
| | | | | | Indel | 0.9461 | 0.6818 | 0.7925 | 58,300 | 3,322 | 27,203 |
| | | | | 0.30 | SNV | 0.9852 | 0.9911 | 0.9882 | 650,570 | 9,773 | 5,824 |
| | | | | | Indel | 0.9436 | 0.8037 | 0.8680 | 69,141 | 4,136 | 16,884 |
| | | DeepSomatic | 50x/25x | 0.05 | SNV | 0.9960 | 0.8560 | 0.9207 | 240,044 | 974 | 40,396 |
| | | | | | Indel | 0.9173 | 0.0237 | 0.0461 | 1,143 | 103 | 47,156 |
| | | | | 0.10 | SNV | 0.9965 | 0.9250 | 0.9594 | 523,877 | 1,837 | 42,470 |
| | | | | | Indel | 0.9901 | 0.1737 | 0.2955 | 13,127 | 131 | 62,465 |
| | | | | 0.20 | SNV | 0.9965 | 0.9604 | 0.9781 | 627,563 | 2,227 | 25,865 |
| | | | | | Indel | 0.9942 | 0.5775 | 0.7306 | 49,375 | 288 | 36,128 |
| | | | | 0.30 | SNV | 0.9963 | 0.9717 | 0.9839 | 637,842 | 2,359 | 18,552 |
| | | | | | Indel | 0.9938 | 0.7396 | 0.8481 | 63,623 | 395 | 22,402 |
| | Single-sample mode | Clair-Mosaic | 50x | 0.05 | SNV | 0.9399 | 0.9643 | 0.9520 | 270,435 | 17,293 | 10,005 |
| | | | | | Indel | 0.3481 | 0.2443 | 0.2871 | 11,799 | 22,096 | 36,500 |
| | | | | 0.10 | SNV | 0.9686 | 0.9826 | 0.9756 | 556,513 | 18,035 | 9,834 |
| | | | | | Indel | 0.6321 | 0.4791 | 0.5450 | 36,213 | 21,077 | 39,379 |
| | | | | 0.20 | SNV | 0.9789 | 0.9789 | 0.9789 | 639,638 | 13,780 | 13,790 |
| | | | | | Indel | 0.8283 | 0.7162 | 0.7682 | 61,236 | 12,698 | 24,267 |
| | | | | 0.30 | SNV | 0.9846 | 0.9827 | 0.9836 | 645,030 | 10,118 | 11,364 |
| | | | | | Indel | 0.9073 | 0.7884 | 0.8437 | 67,826 | 6,926 | 18,199 |
| | | DeepSomatic | 50x | 0.05 | SNV | 0.9066 | 0.1926 | 0.3177 | 54,016 | 5,568 | 226,424 |
| | | | | | Indel | 0.1025 | 0.0113 | 0.0203 | 545 | 4,773 | 47,754 |
| | | | | 0.10 | SNV | 0.9536 | 0.1986 | 0.3287 | 112,458 | 5,466 | 453,889 |
| | | | | | Indel | 0.4745 | 0.0518 | 0.0933 | 3,912 | 4,333 | 71,680 |
| | | | | 0.20 | SNV | 0.9623 | 0.1966 | 0.3265 | 128,463 | 5,034 | 524,965 |
| | | | | | Indel | 0.7135 | 0.1042 | 0.1818 | 8,909 | 3,577 | 76,594 |
| | | | | 0.30 | SNV | 0.9657 | 0.1779 | 0.3005 | 116,775 | 4,147 | 539,619 |
| | | | | | Indel | 0.7408 | 0.0933 | 0.1657 | 8,022 | 2,807 | 78,003 |

**b**

| Dataset | Mode | Caller | Coverage | Sample purity | Variant type | Precision | Recall | F1-score | TP | FP | FN |
|---|---|---|---|---|---|---|---|---|---|---|---|
| PacBio HG002/HG004 mixtures | Paired-sample mode | Clair-Mosaic | 50x/25x | 0.05 | SNV | 0.9687 | 0.9946 | 0.9815 | 280,513 | 9,057 | 1,530 |
| | | | | | Indel | 0.8199 | 0.1811 | 0.2967 | 8,864 | 1,947 | 40,070 |
| | | | | 0.10 | SNV | 0.9814 | 0.9976 | 0.9895 | 558,330 | 10,566 | 1,333 |
| | | | | | Indel | 0.9219 | 0.3814 | 0.5396 | 28,519 | 2,417 | 46,254 |
| | | | | 0.20 | SNV | 0.9805 | 0.9984 | 0.9894 | 641,880 | 12,759 | 1,014 |
| | | | | | Indel | 0.9445 | 0.6684 | 0.7828 | 56,326 | 3,309 | 27,939 |
| | | | | 0.30 | SNV | 0.9839 | 0.9926 | 0.9882 | 641,064 | 10,461 | 4,793 |
| | | | | | Indel | 0.9459 | 0.7969 | 0.8650 | 67,606 | 3,870 | 17,225 |
| | | DeepSomatic | 50x/25x | 0.05 | SNV | 0.9968 | 0.8449 | 0.9146 | 238,292 | 764 | 43,751 |
| | | | | | Indel | 0.9273 | 0.0243 | 0.0473 | 1,187 | 93 | 47,747 |
| | | | | 0.10 | SNV | 0.9961 | 0.9243 | 0.9589 | 517,272 | 2,001 | 42,391 |
| | | | | | Indel | 0.9908 | 0.1812 | 0.3064 | 13,549 | 126 | 61,224 |
| | | | | 0.20 | SNV | 0.9959 | 0.9643 | 0.9798 | 619,927 | 2,566 | 22,967 |
| | | | | | Indel | 0.9946 | 0.5802 | 0.7329 | 48,890 | 265 | 35,375 |
| | | | | 0.30 | SNV | 0.9960 | 0.9719 | 0.9838 | 627,723 | 2,491 | 18,134 |
| | | | | | Indel | 0.9941 | 0.7368 | 0.8463 | 62,502 | 371 | 22,329 |
| | Single-sample mode | Clair-Mosaic | 50x | 0.05 | SNV | 0.9479 | 0.9565 | 0.9522 | 269,777 | 14,833 | 12,266 |
| | | | | | Indel | 0.3443 | 0.2263 | 0.2731 | 11,074 | 21,086 | 37,860 |
| | | | | 0.10 | SNV | 0.9720 | 0.9798 | 0.9759 | 548,378 | 15,825 | 11,285 |
| | | | | | Indel | 0.5852 | 0.4826 | 0.5290 | 36,085 | 25,581 | 38,688 |
| | | | | 0.20 | SNV | 0.9802 | 0.9778 | 0.9790 | 628,637 | 12,707 | 14,257 |
| | | | | | Indel | 0.8258 | 0.7038 | 0.7599 | 59,303 | 12,509 | 24,962 |
| | | | | 0.30 | SNV | 0.9837 | 0.9825 | 0.9831 | 634,584 | 10,487 | 11,273 |
| | | | | | Indel | 0.9062 | 0.7808 | 0.8388 | 66,236 | 6,857 | 18,595 |
| | | DeepSomatic | 50x | 0.05 | SNV | 0.9571 | 0.1863 | 0.3120 | 52,557 | 2,353 | 229,486 |
| | | | | | Indel | 0.1219 | 0.0121 | 0.0220 | 593 | 4,272 | 48,341 |
| | | | | 0.10 | SNV | 0.9764 | 0.1928 | 0.3220 | 107,913 | 2,603 | 451,750 |
| | | | | | Indel | 0.5046 | 0.0522 | 0.0946 | 3,901 | 3,830 | 70,872 |
| | | | | 0.20 | SNV | 0.9790 | 0.1919 | 0.3209 | 123,377 | 2,650 | 519,517 |
| | | | | | Indel | 0.7338 | 0.1040 | 0.1821 | 8,760 | 3,178 | 75,505 |
| | | | | 0.30 | SNV | 0.9781 | 0.1729 | 0.2938 | 111,655 | 2,505 | 534,202 |
| | | | | | Indel | 0.7406 | 0.0914 | 0.1627 | 7,753 | 2,715 | 77,078 |

**Supplementary Table 6. Performance with multiple sample purities in Illumina synthetic datasets.**

Benchmark results of multiple sample purities of different callers using **(a)** HG002/HG003, **(b)** HG002/HG004 mixtures in the Illumina dataset.

**a**

| Dataset | Mode | Caller | Coverage | Sample purity | Variant type | Precision | Recall | F1-score | TP | FP | FN |
|---|---|---|---|---|---|---|---|---|---|---|---|
| Illumina HG002/HG003 mixtures | Paired-sample mode | Clair-Mosaic | 50x/25x | 0.05 | SNV | 0.9844 | 0.7860 | 0.8741 | 256,030 | 4,047 | 69,705 |
| | | | | 0.10 | SNV | 0.9882 | 0.8889 | 0.9359 | 518,376 | 6,189 | 64,774 |
| | | | | 0.20 | SNV | 0.9858 | 0.9647 | 0.9752 | 627,777 | 9,037 | 22,955 |
| | | | | 0.30 | SNV | 0.9865 | 0.9734 | 0.9799 | 635,780 | 8,679 | 17,356 |
| | | DeepSomatic | 50x/25x | 0.05 | SNV | 0.9952 | 0.5392 | 0.6994 | 175,633 | 846 | 150,102 |
| | | | | 0.10 | SNV | 0.9959 | 0.7856 | 0.8783 | 458,136 | 1,899 | 125,014 |
| | | | | 0.20 | SNV | 0.9959 | 0.9448 | 0.9697 | 614,814 | 2,547 | 35,918 |
| | | | | 0.30 | SNV | 0.9962 | 0.9520 | 0.9736 | 621,781 | 2,382 | 31,355 |
| | Single-sample mode | Clair-Mosaic | 50x | 0.05 | SNV | 0.8914 | 0.6372 | 0.7432 | 207,556 | 25,286 | 118,179 |
| | | | | 0.10 | SNV | 0.9511 | 0.8624 | 0.9046 | 502,886 | 25,847 | 80,264 |
| | | | | 0.20 | SNV | 0.9620 | 0.9778 | 0.9698 | 636,276 | 25,144 | 14,456 |
| | | | | 0.30 | SNV | 0.9795 | 0.9693 | 0.9744 | 633,069 | 13,258 | 20,067 |
| | | DeepSomatic | 50x | 0.05 | SNV | 0.7971 | 0.0891 | 0.1602 | 29,011 | 7,386 | 296,724 |
| | | | | 0.10 | SNV | 0.9150 | 0.1373 | 0.2388 | 80,093 | 7,436 | 503,057 |
| | | | | 0.20 | SNV | 0.9426 | 0.1592 | 0.2724 | 103,593 | 6,313 | 547,139 |
| | | | | 0.30 | SNV | 0.9563 | 0.1535 | 0.2645 | 100,256 | 4,577 | 552,880 |

**b**

| Dataset | Mode | Caller | Coverage | Sample purity | Variant type | Precision | Recall | F1-score | TP | FP | FN |
|---|---|---|---|---|---|---|---|---|---|---|---|
| Illumina HG002/HG004 mixtures | Paired-sample mode | Clair-Mosaic | 50x/25x | 0.05 | SNV | 0.9834 | 0.8030 | 0.8841 | 258,887 | 4,360 | 63,512 |
| | | | | 0.10 | SNV | 0.9882 | 0.8996 | 0.9418 | 517,712 | 6,173 | 57,774 |
| | | | | 0.20 | SNV | 0.9854 | 0.9687 | 0.9770 | 620,538 | 9,223 | 20,020 |
| | | | | 0.30 | SNV | 0.9862 | 0.9756 | 0.9809 | 627,312 | 8,799 | 15,668 |
| | | DeepSomatic | 50x/25x | 0.05 | SNV | 0.9947 | 0.5483 | 0.7069 | 176,771 | 938 | 145,628 |
| | | | | 0.10 | SNV | 0.9955 | 0.7882 | 0.8798 | 453,575 | 2,028 | 121,911 |
| | | | | 0.20 | SNV | 0.9955 | 0.9485 | 0.9714 | 607,585 | 2,747 | 32,973 |
| | | | | 0.30 | SNV | 0.9957 | 0.9558 | 0.9753 | 614,554 | 2,656 | 28,426 |
| | Single-sample mode | Clair-Mosaic | 50x | 0.05 | SNV | 0.8849 | 0.6483 | 0.7484 | 209,025 | 27,191 | 113,374 |
| | | | | 0.10 | SNV | 0.9476 | 0.8682 | 0.9062 | 499,642 | 27,644 | 75,844 |
| | | | | 0.20 | SNV | 0.9599 | 0.9792 | 0.9694 | 627,250 | 26,228 | 13,308 |
| | | | | 0.30 | SNV | 0.9715 | 0.9808 | 0.9762 | 630,666 | 18,503 | 12,314 |
| | | DeepSomatic | 50x | 0.05 | SNV | 0.8009 | 0.0899 | 0.1617 | 28,989 | 7,206 | 293,410 |
| | | | | 0.10 | SNV | 0.9142 | 0.1352 | 0.2355 | 77,800 | 7,301 | 497,686 |
| | | | | 0.20 | SNV | 0.9369 | 0.1568 | 0.2687 | 100,454 | 6,760 | 540,104 |
| | | | | 0.30 | SNV | 0.9576 | 0.1508 | 0.2606 | 96,959 | 4,293 | 546,021 |

**Supplementary Table 7. Performance on the HG002 real dataset in different platforms.**

Benchmark results on HG002 dataset using **(a)** ONT, **(b)** PacBio, **(c)** Illumina platforms.

**a**

| Dataset | Mode | Caller | Coverage | Variant type | Precision | Recall | F1-score | TP | FP | FN |
|---|---|---|---|---|---|---|---|---|---|---|
| ONT HG002 | Paired-sample mode | Clair-Mosaic | 50x/50x | SNV | 0.0414 | 0.9872 | 0.0795 | 77 | 1,782 | 1 |
| | | DeepSomatic | 50x/50x | SNV | 0.0007 | 0.9744 | 0.0013 | 76 | 115,768 | 2 |
| | Single-sample mode | Clair-Mosaic | 50x | SNV | 0.0039 | 0.9487 | 0.0078 | 74 | 19,119 | 4 |
| | | DeepSomatic | 50x | SNV | 0.0003 | 0.9231 | 0.0007 | 72 | 212,484 | 6 |

**b**

| Dataset | Mode | Caller | Coverage | Variant type | Precision | Recall | F1-score | TP | FP | FN |
|---|---|---|---|---|---|---|---|---|---|---|
| PacBio HG002 | Paired-sample mode | Clair-Mosaic | 50x/50x | SNV | 0.1258 | 0.9762 | 0.2229 | 82 | 570 | 2 |
| | | DeepSomatic | 50x/50x | SNV | 0.0032 | 0.9643 | 0.0063 | 81 | 25,372 | 3 |
| | Single-sample mode | Clair-Mosaic | 50x | SNV | 0.0056 | 0.9643 | 0.0111 | 81 | 14,306 | 3 |
| | | DeepSomatic | 50x | SNV | 0.0004 | 0.9405 | 0.0008 | 79 | 190,604 | 5 |

**c**

| Dataset | Mode | Caller | Coverage | Variant type | Precision | Recall | F1-score | TP | FP | FN |
|---|---|---|---|---|---|---|---|---|---|---|
| Illumina HG002 | Paired-sample mode | Clair-Mosaic | 50x/50x | SNV | 0.0080 | 0.9747 | 0.0159 | 77 | 9,490 | 2 |
| | | DeepSomatic | 50x/50x | SNV | 0.0001 | 0.9367 | 0.0003 | 74 | 565,886 | 5 |
| | Single-sample mode | Clair-Mosaic | 50x | SNV | 0.0041 | 0.9367 | 0.0082 | 74 | 18,045 | 5 |
| | | DeepSomatic | 50x | SNV | 0.0004 | 0.9241 | 0.0008 | 73 | 178,194 | 6 |
| | | MosaicHunter | 50x | SNV | 0.0005 | 0.7468 | 0.0010 | 75 | 180,542 | 4 |
| | | MosaicForecast | 50x | SNV | 0.1574 | 0.8228 | 0.2642 | 75 | 48,033 | 4 |
| | | DeepMosaic | 50x | SNV | 0.0191 | 0.9367 | 0.0375 | 59 | 114,520 | 20 |

### Supplementary Methods

#### Pileup input

The pileup input of Clair-Mosaic comprises 1,122 integers (i.e., 33 positions with 34 features at each position). The 34 features provide information on the counts on both the forward and reverse strand of 1) nucleotides, 2) insertions, 3) deletions, 4) nucleotides with low mapping quality, and 5) nucleotides with low base quality. Below is a detailed breakdown of each feature:

1-4: A+/C+/G+/T+: The counts of A/C/G/T nucleotides present in the forward strand.

5: $I_S$+: The count of insertions that have the same starting positions as the candidate site in the forward strand.

6: $I_{1S}$+: Similar to $I_S$+ but only counting the insertion with the highest read support.

7: $D_S$+: The count of deletions that have the same starting positions as the candidate site in the forward strand.

8: $D_{1S}$+: Similar to $D_S$+ but only counting the deletion with the highest read support.

9: $D_R$+: The count of all deletions in the forward strand that crossed the position except for the first base of each deletion.

10-13: A-/C-/G-/T-: The counts of A/C/G/T nucleotides present in the reverse strand.

14: $I_S$-: The count of insertions that have the same starting positions as the candidate site in the reverse strand.

15: $I_{1S}$-: Similar to $I_S$- but only counting the insertion with the highest read support.

16: $D_S$-: The count of deletions that have the same starting positions as the candidate site in the reverse strand.

17: $D_{1S}$-: Similar to $D_S$- but only counting the deletion with the highest read support.

18: $D_R$-: The count of all deletions in the reverse strand that crossed the position except for the first base of each deletion.

19-22: $A_{LMQ}$+/$C_{LMQ}$+/$G_{LMQ}$+/$T_{LMQ}$+: The counts of A/C/G/T nucleotides with low mapping quality (MQ<20) in the forward strand.

23-26: $A_{LBQ}$+/$C_{LBQ}$+/$G_{LBQ}$+/$T_{LBQ}$+: The counts of A/C/G/T nucleotides with low base quality (BQ<30) in the forward strand.

27-30: $A_{LMQ}$-/$C_{LMQ}$-/$G_{LMQ}$-/$T_{LMQ}$-: The counts of A/C/G/T nucleotides with low mapping quality (MQ<20) in the reverse strand.

31-34: $A_{LBQ}$-/$C_{LBQ}$-/$G_{LBQ}$-/$T_{LBQ}$-: The counts of A/C/G/T nucleotides with low base quality (BQ<30) in the reverse strand.

### Command lines used

#### Read alignment

##### Minimap2 (v2.17-r941)

```
# Align ONT reads using minimap2 to GRCh38_no_alt by default
minimap2 -t ${THREADS} -aL -z 600,200 -x map-ont ref.fa input.fastq.gz | samtools view -bh
-o output.unsorted.bam -
samtools sort -@${THREADS} -o output.sorted.bam output.unsorted.bam && samtools index -@
${THREADS} output.sorted.bam
```

#### BAM subsampling

##### Samtools (v1.15.1)

```
samtools view -@ ${THREADS} -s ${RATIO}.${RATIO} -b -o subsampled.bam ${BAM}
samtools index -@ ${THREADS} subsampled.bam
```

#### Coverage calculation

##### Mosdepth (v0.3.1)

```
mosdepth -t ${THREADS} -n -x --quantize 0:15:150: output ${BAM}
```

#### Alignment statistical summary

##### NanoPlot (v1.40.2)

```
NanoPlot -t ${THREADS} --bam ${BAM} --N50 -o bamplots
```

#### Generating BAMs with different sample purities

```
# Use ${SAMPLE_PURITY} to add sample purity
pypy3 clair_mosaic.py gen_contaminated_bam \
       --sample_bam_fn ${SAMPLE_BAM_FILE_PATH} \
       --control_bam_fn ${CONTROL_BAM_FILE_PATH} \
       --sample_purity ${SAMPLE_PURITY} \
       --output_dir ${OUTPUT_DIR} \
       --sample_bam_coverage ${SAMPLE_BAM_COVERAGE} \
       --control_bam_coverage ${CONTROL_BAM_COVERAGE} \
       --mosdepth ${MOSDEPTH_PATH}
```

#### Running Clair-Mosaic for ONT data

##### Clair-Mosaic (v0.0.1)

```
# For paired-sample mode
docker run -it \
```

```
  -v ${INPUT_DIR}:${INPUT_DIR} \
  -v ${OUTPUT_DIR}:${OUTPUT_DIR} \
  hkubal/clair-mosaic:v0.0.1 \
  /opt/bin/run_clair_mosaic \
       --bam_fn ${BAM_FILE_PATH} \
       --control_bam_fn ${CONTROL_BAM_FILE_PATH} \
       --ref_fn ${REF} \
       --threads ${THREADS} \
       --platform ont \
       --output_dir ${OUTPUT_DIR}

# For single-sample mode
docker run -it \
  -v ${INPUT_DIR}:${INPUT_DIR} \
  -v ${OUTPUT_DIR}:${OUTPUT_DIR} \
  hkubal/clair-mosaic:v0.0.1 \
  /opt/bin/run_clair_mosaic \
       --bam_fn ${BAM_FILE_PATH} \
       --ref_fn ${REF} \
       --threads ${THREADS} \
       --platform ont \
       --output_dir ${OUTPUT_DIR}
```

#### Running other mosaic variant callers for ONT data

##### DeepSomatic (v1.7.0)

```
# For paired-sample mode
docker run -it \
  -v ${INPUT_DIR}:${INPUT_DIR} \
  -v ${OUTPUT_DIR}:${OUTPUT_DIR} \
  google/deepsomatic:v17_rc1_08082024 \
  run_deepsomatic \
       --model_type="ONT" \
       --ref=${REF} \
       --reads_tumor=${TUMOR_BAM_FILE_PATH} \
       --reads_normal=${NORMAL_BAM_FILE_PATH} \
       --output_vcf=${OUTPUT_DIR}/output.vcf.gz \
       --sample_name_tumor="tumor" \
       --sample_name_normal="normal" \
       --num_shards=${THREADS}

# For single-sample mode
docker run -it \
  -v ${INPUT_DIR}:${INPUT_DIR} \
```

```
  -v ${OUTPUT_DIR}:${OUTPUT_DIR} \
  google/deepsomatic:v17_rc1_08082024 \
  run_deepsomatic \
        --model_type="ONT_TUMOR_ONLY" \
        --ref=${REF} \
        --reads_tumor=${TUMOR_BAM_FILE_PATH} \
        --output_vcf=${OUTPUT_DIR}/output.vcf.gz \
        --sample_name_tumor="tumor" \
        --num_shards=${THREADS} \
        --use_default_pon_filtering True
```

#### Running Clair-Mosaic for PacBio data

##### Clair-Mosaic (v0.0.1)

```
# For paired-sample mode
docker run -it \
  -v ${INPUT_DIR}:${INPUT_DIR} \
  -v ${OUTPUT_DIR}:${OUTPUT_DIR} \
  hkubal/clair-mosaic:v0.0.1 \
  /opt/bin/run_clair_mosaic \
        --bam_fn ${BAM_FILE_PATH} \
        --control_bam_fn ${CONTROL_BAM_FILE_PATH} \
        --ref_fn ${REF} \
        --threads ${THREADS} \
        --platform hifi_revio \
        --output_dir ${OUTPUT_DIR}

# For single-sample mode
docker run -it \
  -v ${INPUT_DIR}:${INPUT_DIR} \
  -v ${OUTPUT_DIR}:${OUTPUT_DIR} \
  hkubal/clair-mosaic:v0.0.1 \
  /opt/bin/run_clair_mosaic \
        --bam_fn ${BAM_FILE_PATH} \
        --ref_fn ${REF} \
        --threads ${THREADS} \
        --platform hifi_revio \
        --output_dir ${OUTPUT_DIR}
```

#### Running other mosaic variant callers for PacBio data

##### DeepSomatic (v1.7.0)

```
# For paired-sample mode
docker run -it \
```

```
  -v ${INPUT_DIR}:${INPUT_DIR} \
  -v ${OUTPUT_DIR}:${OUTPUT_DIR} \
  google/deepsomatic:v17_rc1_08082024 \
  run_deepsomatic \
       --model_type="PACBIO" \
       --ref=${REF} \
       --reads_tumor=${TUMOR_BAM_FILE_PATH} \
       --reads_normal=${NORMAL_BAM_FILE_PATH} \
       --output_vcf=${OUTPUT_DIR}/output.vcf.gz \
       --sample_name_tumor="tumor" \
       --sample_name_normal="normal" \
       --num_shards=${THREADS}

# For single-sample mode
docker run -it \
  -v ${INPUT_DIR}:${INPUT_DIR} \
  -v ${OUTPUT_DIR}:${OUTPUT_DIR} \
  google/deepsomatic:v17_rc1_08082024 \
  run_deepsomatic \
       --model_type="PACBIO_TUMOR_ONLY" \
       --ref=${REF} \
       --reads_tumor=${TUMOR_BAM_FILE_PATH} \
       --output_vcf=${OUTPUT_DIR}/output.vcf.gz \
       --sample_name_tumor="tumor" \
       --num_shards=${THREADS} \
       --use_default_pon_filtering True
```

#### Running Clair-Mosaic for Illumina data

##### Clair-Mosaic (v0.0.1)

```
# For paired-sample mode
docker run -it \
  -v ${INPUT_DIR}:${INPUT_DIR} \
  -v ${OUTPUT_DIR}:${OUTPUT_DIR} \
  hkubal/clair-mosaic:v0.0.1 \
  /opt/bin/run_clair_mosaic \
       --bam_fn ${BAM_FILE_PATH} \
       --control_bam_fn ${CONTROL_BAM_FILE_PATH} \
       --ref_fn ${REF} \
       --threads ${THREADS} \
       --platform ilmn \
       --output_dir ${OUTPUT_DIR}

# For single-sample mode
```

```
docker run -it \
  -v ${INPUT_DIR}:${INPUT_DIR} \
  -v ${OUTPUT_DIR}:${OUTPUT_DIR} \
  hkubal/clair-mosaic:v0.0.1 \
  /opt/bin/run_clair_mosaic \
       --bam_fn ${BAM_FILE_PATH} \
       --ref_fn ${REF} \
       --threads ${THREADS} \
       --platform ilmn \
       --output_dir ${OUTPUT_DIR}
```

#### Running other mosaic variant callers for Illumina data

##### DeepSomatic (v1.7.0)

```
# For paired-sample mode
docker run -it \
  -v ${INPUT_DIR}:${INPUT_DIR} \
  -v ${OUTPUT_DIR}:${OUTPUT_DIR} \
  google/deepsomatic:v17_rc1_08082024 \
  run_deepsomatic \
       --model_type="WGS" \
       --ref=${REF} \
       --reads_tumor=${TUMOR_BAM_FILE_PATH} \
       --reads_normal=${NORMAL_BAM_FILE_PATH} \
       --output_vcf=${OUTPUT_DIR}/output.vcf.gz \
       --sample_name_tumor="tumor" \
       --sample_name_normal="normal" \
       --num_shards=${THREADS}

# For single-sample mode
docker run -it \
  -v ${INPUT_DIR}:${INPUT_DIR} \
  -v ${OUTPUT_DIR}:${OUTPUT_DIR} \
  google/deepsomatic:v17_rc1_08082024 \
  run_deepsomatic \
       --model_type="WGS_TUMOR_ONLY" \
       --ref=${REF} \
       --reads_tumor=${TUMOR_BAM_FILE_PATH} \
       --output_vcf=${OUTPUT_DIR}/output.vcf.gz \
       --sample_name_tumor="tumor" \
       --num_shards=${THREADS} \
       --use_default_pon_filtering True
```

##### MosaicHunter (v1.0.0)

```
java -jar build/mosaichunter.jar genome \
       -P input_file=${BAM_FILE_PATH} \
       -P reference_file=${REF} \
       -P output_dir=${OUTPUT_DIR} \
       -P valid_references=${CHR}
```

#### MosaicForecast (v0.0.1)

```
docker run -it \
  -v ${INPUT_DIR}:${INPUT_DIR} \
  -v ${OUTPUT_DIR}:${OUTPUT_DIR} \
  yanmei/mosaicforecast:0.0.1 \
  python ReadLevel_Features_extraction.py \
       ${SAMPLE_INPUT} \
       ${SAMPLE_FEATURES} \
       ${BAM_DIR} \
       ${REF} \
       ${UMAP_PATH} \
       ${THREADS} \
       bam

docker run -it \
  -v ${INPUT_DIR}:${INPUT_DIR} \
  -v ${OUTPUT_DIR}:${OUTPUT_DIR} \
  yanmei/mosaicforecast:0.0.1 \
  Rscript Prediction.R \
       ${SAMPLE_FEATURES} \
       ${MODEL_PATH} \
       Refine \
       ${OUTPUT}
```

#### DeepMosaic (v1.1.4)

```
singularity exec \
  -B ${INPUT_DIR}:${INPUT_DIR} \
  -B ${OUTPUT_DIR}:${OUTPUT_DIR} \
  deepmosaic.sif \
  deepmosaic-draw \
       -i ${SAMPLE_INPUT} \
       -o ${OUTPUT_DIR} \
       -a annovar \
       -b hg38 \
       -db gnomad_genome

singularity exec \
```

```
  -B ${INPUT_DIR}:${INPUT_DIR} \
  -B ${OUTPUT_DIR}:${OUTPUT_DIR} \
  deepmosaic.sif \
  deepmosaic-predict \
       -i ${SAMPLE_FEATURES} \
       -o ${OUTPUT} \
       -m ${MODEL_PATH} \
       -b ${THREADS} \
       -gb hg38
```

#### Benchmarking

Calculate Precision, Recall, F1-Score with "compare_vcf" submodule in Clair-Mosaic

```
pypy3 clair_mosaic.py compare_vcf \
       --truth_vcf_fn ${TRUTH_VCF} \
       --input_vcf_fn output.vcf.gz \
       --bed_fn ${TRUTH_CONFIDENT_BED} \
       --output_dir benchmark_result \
       --input_filter_tag 'PASS' \
       --min_qual ${MIN_QUAL} \
       --min_af ${MIN_AF} \
       --output_best_f1_score \
       --bam_fn ${TUMOR_BAM_FILE_PATH}
```

### Data availability

#### GIAB truth variants

HG001 (NA12878), GRCh38, v4.2.1
https://ftp-trace.ncbi.nlm.nih.gov/giab/ftp/release/NA12878_HG001/NISTv4.2.1/GRCh38/

HG002 (NA24385), GRCh38, v4.2.1
https://ftp-trace.ncbi.nlm.nih.gov/giab/ftp/release/AshkenazimTrio/HG002_NA24385_son/NISTv4.2.1/GRCh38/

HG003 (NA24149), GRCh38, v4.2.1
https://ftp-trace.ncbi.nlm.nih.gov/giab/ftp/release/AshkenazimTrio/HG003_NA24149_father/NISTv4.2.1/GRCh38/

HG004 (NA24143), GRCh38, v4.2.1
https://ftp-trace.ncbi.nlm.nih.gov/giab/ftp/release/AshkenazimTrio/HG004_NA24143_mother/NISTv4.2.1/GRCh38/

HG005 (NA24631), GRCh38, v4.2.1
https://ftp-trace.ncbi.nlm.nih.gov/giab/ftp/release/ChineseTrio/HG005_NA24631_son/NISTv4.2.1/GRCh38/

HG006 (NA24694), GRCh38, v4.2.1
https://ftp-trace.ncbi.nlm.nih.gov/giab/ftp/release/ChineseTrio/HG006_NA24694_father/NISTv4.2.1/GRCh38/

HG007 (NA24695), GRCh38, v4.2.1

https://ftp-trace.ncbi.nlm.nih.gov/giab/ftp/release/ChineseTrio/HG007_NA24695_mother/NISTv4.2.1/GRCh38/

HG002 mosaic variant benchmark, GRCh38, v1.0
https://ftp-trace.ncbi.nlm.nih.gov/giab/ftp/release/AshkenazimTrio/HG002_NA24385_son/mosaic_v1.00/GRCh38/SNV/

#### Reference genomes

GRCh38_no_alt
https://ftp.ncbi.nlm.nih.gov/genomes/all/GCA/000/001/405/GCA_000001405.15_GRCh38/seqs_for_alignment_pipelines.ucsc_ids/GCA_000001405.15_GRCh38_no_alt_analysis_set.fna.gz

GRCh38 d1.vd1
https://ftp-trace.ncbi.nlm.nih.gov/ReferenceSamples/seqc/Somatic_Mutation_WG/technical/reference_genome/GRCh38/GRCh38.d1.vd1.fa

GRCh38 Stratification regions (v3.3)
https://ftp-trace.ncbi.nlm.nih.gov/giab/ftp/release/genome-stratifications/v3.3/GRCh38@all

#### ONT Sequencing Data

ONT EPI2ME Labs HG001 R10.4.1 Q20+, GRCh38_no_alt, 78.89-fold
https://labs.epi2me.io/giab-2023.05/

ONT EPI2ME Labs HG002 R10.4.1 Q20+, GRCh38_no_alt, 91.18-fold
https://labs.epi2me.io/giab-2023.05/

ONT EPI2ME Labs HG003 R10.4.1 Q20+, GRCh38_no_alt, 72.51-fold
https://labs.epi2me.io/giab-2023.05/

ONT EPI2ME Labs HG004 R10.4.1 Q20+, GRCh38_no_alt, 58.84-fold
https://labs.epi2me.io/giab-2023.05/

ONT EPI2ME Labs HG005 R10.4.1 Q20+, GRCh38_no_alt, 55.00-fold
https://labs.epi2me.io/giab-2025.01/

ONT EPI2ME Labs HG006 R10.4.1 Q20+, GRCh38_no_alt, 54.89-fold
https://labs.epi2me.io/giab-2025.01/

ONT EPI2ME Labs HG007 R10.4.1 Q20+, GRCh38_no_alt, 65.95-fold
https://labs.epi2me.io/giab-2025.01/

#### PacBio Sequencing Data

PacBio HG002 HiFi Revio, GRCh38_no_alt, 90.28-fold
https://downloads.pacbcloud.com/public/revio/2022Q4/HG002-rep1/

PacBio HG003 HiFi Revio, GRCh38_no_alt, 61.15-fold
https://downloads.pacbcloud.com/public/revio/2022Q4/HG003-rep1/

PacBio HG004 HiFi Revio, GRCh38_no_alt, 62.67-fold
https://downloads.pacbcloud.com/public/revio/2022Q4/HG004-rep1/

#### Illumina Sequencing Data

HG002 NovaSeq 6000 (NA24385), GRCh38_no_alt, 49.82-fold
https://storage.googleapis.com/brain-genomics-public/research/sequencing/grch38/bam/novaseq/wgs_pcr_free/50x/HG002.novaseq.pcr-free.50x.dedup.grch38.bam

HG003 NovaSeq 6000 (NA24149), GRCh38_no_alt, 47.38-fold
https://storage.googleapis.com/brain-genomics-public/research/sequencing/grch38/bam/novaseq/wgs_pcr_free/50x/HG003.novaseq.pcr-free.50x.dedup.grch38.bam

HG004 NovaSeq 6000 (NA24143), GRCh38_no_alt, 46.36-fold
https://storage.googleapis.com/brain-genomics-public/research/sequencing/grch38/bam/novaseq/wgs_pcr_free/50x/HG004.novaseq.pcr-free.50x.dedup.grch38.bam

HG003 HiSeqX (NA24149), GRCh38_no_alt 43.77-fold
https://storage.googleapis.com/brain-genomics-public/research/sequencing/grch38/bam/hiseqx/wgs_pcr_free/40x/HG003.hiseqx.pcr-free.40x.dedup.grch38.bam

HG004 HiSeqX (NA24143), GRCh38_no_alt, 42.13-fold
https://storage.googleapis.com/brain-genomics-public/research/sequencing/grch38/bam/hiseqx/wgs_pcr_free/40x/HG004.hiseqx.pcr-free.40x.dedup.grch38.bam
